# From sequence to signature: Machine learning uncovers multiscale feature landscapes that predict AMR across ESKAPE pathogens

**DOI:** 10.1101/2025.07.03.663053

**Authors:** Abhirupa Ghosh, Evan P Brenner, Charmie K Vang, Ethan P Wolfe, Emily A Boyer, Raymond L Lesiyon, Keenan R Manpearl, Vignesh Sridhar, Joseph T Burke, Jacob D Krol, Jill MR Bilodeaux, Janani Ravi

## Abstract

Since the clinical introduction of antibiotics in the 1940s, antimicrobial resistance (AMR) has become an increasingly dire threat to global public health. Pathogens acquire AMR much faster than we discover new drugs (antibiotics), warranting innovative methods to better understand its molecular underpinnings. Traditional approaches for detecting AMR in novel bacterial strains are time-consuming and labor-intensive. However, advances in sequencing technology offer a plethora of bacterial genome data, and computational approaches like machine learning (ML) provide an optimistic scope for in silico AMR prediction. Here, we introduce a comprehensive multiscale ML approach to predict AMR phenotypes and identify AMR molecular features associated with a single drug or drug family, stratified by time and geographical locations. As a case study, we focus on a subset of the World Health Organization’s Bacterial Priority Pathogens, the frequently drug-resistant and nosocomial ESKAPE pathogens: Enterococcus faecium, Staphylococcus aureus, Klebsiella pneumoniae, Acinetobacter baumannii, Pseudomonas aeruginosa, and Enterobacter species. We started with sequenced genomes with lab-derived AMR phenotypes, constructed pangenomes, clustered gene and protein sequences, and extracted protein domains to generate pangenomic features across molecular scales. To uncover the molecular mechanisms behind drug-/drug class-specific resistance, we trained logistic regression ML models on our datasets. These yielded ranked lists of AMR-associated genes, proteins, and domains. In addition to recapitulating known AMR features, our models identified novel candidates for experimental validation. The models were performant across molecular scales, data types, and drugs while achieving a median normalized Matthews correlation coefficient of 0.89. Prediction performance showed resilience even when evaluated on geographical and temporal holdouts. We also evaluated model generalizability and cross-resistance across the drug-/drug class-specific models cross-tested on other available drug-/drug class genomes. Finally, we uncovered multiple drug class resistance features using multiclass and multilabel models. Our holistic approach promises reliable prediction of existing and developing resistance in newly sequenced pathogen genomes, while pinpointing the mechanistic molecular contributors of AMR. All our models and results are available at our interactive web app, https://jravilab.org/amr.

## Introduction

Antimicrobial resistance (AMR) is a growing global health crisis. The rise of resistant bacterial pathogens is steadily compromising the effectiveness of antibiotics, long regarded as a cornerstone of modern medicine since their clinical integration in the 1940s^1–3^. In the absence of effective therapies, even routine infections and common medical interventions could result in outcomes reminiscent of the pre-antibiotic era (e.g., amputation or quarantine)^1^. Bacterial AMR can be directly attributable to more than one million deaths annually and associated with nearly five million, with the highest burden falling on low-resource settings^4^. The UK Review on Antimicrobial Resistance warns that the direct death toll could reach ten million per year by 2050^5^. Further, the rise of multidrug-resistant (MDR) bacteria, defined as pathogens resistant to three or more classes of antibiotics, has dramatically narrowed viable treatment options^6^. Compounding this threat, resistance to “last-resort” antibiotics such as carbapenems and polymyxins is becoming more common^7,8^. These pan-resistant strains leave clinicians with no effective treatment options, leading to higher mortality rates and impairing outbreak containment. At the same time, there has been a marked decline in the successful development and clinical deployment of new antibiotics^9^. This stagnation not only lags behind the pace of resistance evolution but also is fueled by economic disincentives and regulatory hurdles that have eroded industry investment in antibiotic innovation^10^. Diagnostic limitations further compound the problem since many healthcare settings lack the capacity to identify resistance phenotypes rapidly and accurately, delaying appropriate treatment and allowing resistant strains to spread unchecked. A group of pathogens collectively referred to as ESKAPE (*Enterococcus faecium, Staphylococcus aureus, Klebsiella pneumoniae, Acinetobacter baumannii, Pseudomonas aeruginosa,* and *Enterobacter* species) is among the leading contributors of antibiotic-resistant infections and AMR-associated mortality^11^. The persistence and virulence of these pathogens stem not only from spontaneous mutations and horizontal gene transfer (HGT), but also from their arsenal of intrinsic resistance mechanisms (e.g., biofilm formation) that make them inherently difficult to treat^12^. In clinical settings, this biological advantage is enhanced by human behavior; up to half of antibiotic prescriptions involve unnecessary or incorrect antibiotics^13,14^. This creates sustained selective pressures that propagate AMR emergence and harm patient outcomes.

Although AMR continues to rise, the molecular mechanisms underlying drug resistance remain poorly understood^15–17^. Most drug susceptibility testing only provides a phenotypic readout (i.e., a growth defect in the presence of an antibiotic) but does not reveal *why* resistance occurs^13^. While the genetic determinants have been identified for select drug-organism pairs through rigorous laboratory testing^7,17^, much of the resistome remains unexplored, especially for MDR strains and non-model pathogens. Whole genome sequencing (WGS), when paired with modern technologies like next-generation sequencing^18^, offers unprecedented access to complete bacterial genomes. Over the past decade, the rapid expansion of sequencing capacity and plummeting costs have generated vast genomic datasets^19^. Yet, the integration of WGS into routine AMR surveillance or diagnostics has been limited^13^.

Machine learning (ML) has been applied in recent years to leverage omics datasets for the identification of biomarkers of human disease, such as cancer and heart disease^20^. Researchers have begun exploring the prediction of AMR with ML and deep learning (DL) as well^18,19^. For instance, DeepARG uses a DL framework trained on curated antibiotic resistance gene (ARG) databases to classify short sequencing reads into resistance categories with high precision^21^. Other efforts like AMRLearn use single nucleotide polymorphism (SNP) genotypes in various ML classifiers to predict AMR^22^. Despite their promise, these models face significant limitations. Most rely on known resistance genes catalogued in curated databases (e.g., Comprehensive Antibiotic Resistance Database (CARD^23^), ResFinder^24^), which biases predictions toward previously characterized mechanisms and limits the discovery of novel or rare variants.

Herein, we demonstrate the use of interpretable logistic regression (LR) models to predict AMR across multiple molecular feature scales and species using public data from the Bacterial and Viral Bioinformatics Resource Center (BV-BRC)^25^. By incorporating gene-, protein-, and domain-level features, our models capture distinct layers of biological function, enabling broader generalization and deeper mechanistic insight into resistance determinants. We demonstrate the utility of this workflow by exploring AMR in the context of the ESKAPE pathogens. We chose complete and draft (WGS) genomes with available lab-derived AMR phenotypic data to create feature matrices using the presence/absence (binary) or counts of genes, proteins, domains, and gene neighborhood rearrangements from species-specific pangenomes. We trained and tested models on drugs across multiple classes per species. These models perform well across pathogens, drugs, and feature scales by standard ML performance metrics, provide a testable output of features predicted to be associated with AMR, and are computationally fast and lightweight. An inherent strength of ML is that it does not rely on *a priori* loci annotated to be involved. Instead, our models identify associations between features and genome resistance/susceptibility labels, permitting detection of previously unknown AMR-associated molecular mechanisms. Such approaches are essential as resistance develops and spreads, and new pathogens emerge. This work offers multiscale ML as a holistic method of AMR detection and attribution.

Our framework highlights previously obscure AMR mechanisms and enables the development of nuanced questions. Will one type of molecular feature prove better than another at predicting resistance for a given combination of pathogen and antibiotic? If so, why might that be? How does the format of input data, such as binary (presence/absence of features) vs. counts, impact model performance and interpretability? Which features contribute to resistance across species and drugs? Finally, we leverage temporal and geographic metadata to test the robustness of our models with various holdouts and assess limitations. This allows us to validate the generalizability of the models to genomes from different continents and time periods. Alongside, we identify features that are unique or shared contributing factors across holdouts.

## Methods

### Data collection, preprocessing, and metadata curation

To predict whether a bacterial isolate is resistant or susceptible to a given drug based on its genome and to identify the genomic features that drive these predictions, we developed a computational workflow integrating comparative genomics and supervised ML, as illustrated in **Figure 1**. We downloaded assembled genome sequences, encoded protein sequences, and corresponding isolate metadata, including laboratory-derived drug susceptibility phenotypes per genome, from the Bacterial and Viral Bioinformatics Resource Center (BV-BRC^25^; accessed June 20, 2025) using their BV-BRC command-line interface P3-scripts (`v5.3`) and custom R functions. We focused on a subset of the WHO Bacterial Priority Pathogens List: *Enterococcus faecium* (Efa), *Staphylococcus aureus* (Sau), *Klebsiella pneumoniae* (Kpn)*, Acinetobacter baumannii* (Aba), *Pseudomonas aeruginosa* (Pae), and *Enterobacter cloacae* and *hormaechei*^26^ (*Enterobacter* spp., Esp.), collectively referred as ESKAPE pathogens. Query parameters included both taxon IDs and species names to enable species-specific targeting of divergent and taxonomically complex clades. Genomes were filtered to include only “Good” quality, “Complete” and “Draft/WGS” assemblies, with plasmid sequences when available. Retrieved genome metadata were parsed and stored in local DuckDB^27^ databases in Apache Parquet format to support fast querying, smaller file sizes, and reproducible updates. AMR-associated metadata were downloaded and merged with genome metadata using genome IDs. Reported AMR phenotypes for each genome-drug pair were mapped to binary resistant or susceptible labels. Intermediate phenotypes were excluded due to their low frequency and unclear biological significance in distinguishing true resistance from susceptibility. Drug names were harmonized and mapped to antibiotic classes based on WHO AWaRe guidelines^28^ and AntibioticDB^29^. Metadata curation also included standardizing isolate identifiers and ensuring consistency across genome-, protein-, and domain-level feature matrices.

**Figure 1.**
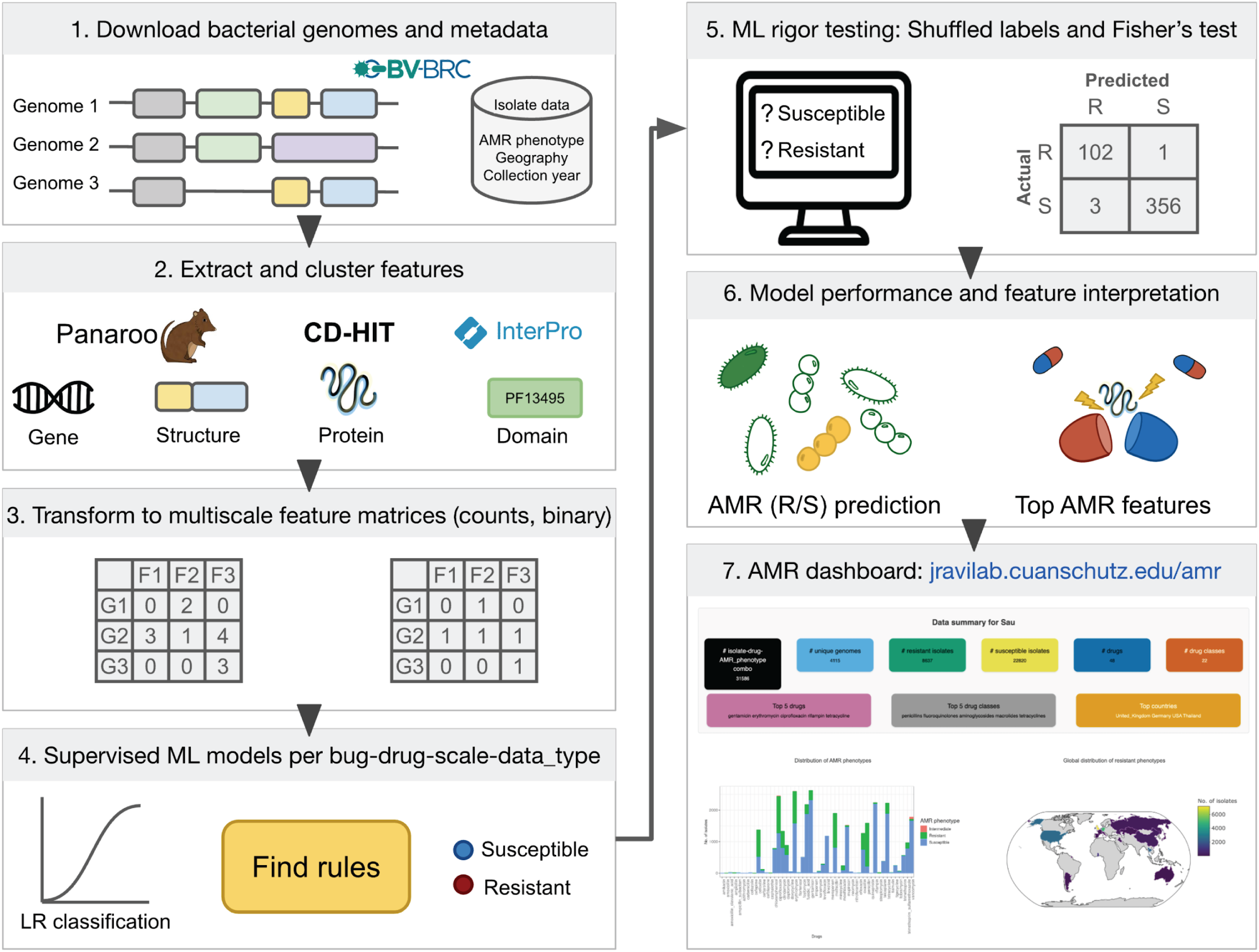
Our workflow to predict AMR from ESKAPE pathogen genomes. We developed a bacterial species-agnostic workflow using public data and supervised machine learning (ML) methods to predict bacterial isolate antimicrobial resistance (AMR) phenotypes against specific drugs. In brief, the pipeline starts with **1)** using the Bacterial and Viral Bioinformatics Resource Center (BV-BRC) to acquire genomic sequences (.fna) and annotated protein sequences (.faa) with paired AMR phenotype labels and metadata from chosen bacterial species. **2)** Features are clustered using Panaroo (genes and gene structure) and CD-HIT (proteins), and domains are extracted from the proteins using InterPro. **3)** Per-genome count and presence/absence data type matrices are created for ML input. **4)** Supervised logistic regression (LR) ML models are trained per species/drug/feature scale/data type on drug resistance/susceptibility labels. **5)** As a baseline, labels are shuffled per matrix and ML models are trained to create a control without biological signals, and non-ML analysis is performed by Fisher’s exact testing of features. **6)** Model performance is compared across species, drugs, molecular scales, and data types, and the top predictive molecular features are analyzed for their relevance to AMR. **7)** We share our results in an interactive data dashboard (jravilab.org/amr) for exploration, visualization, and hypothesis generation.

### Generating pan-feature matrices

*Genes*. We used Panaroo^30^ (`v1.5.2`) to cluster homologous genes and construct bacterial pangenomes, producing both genome-by-gene count matrices (`gene_count`) and presence/absence (binary 1/0) matrices (`gene_binary`) with associated AMR phenotype labels per genome. Genes were grouped based on a 95% sequence identity threshold, allowing up to 5% length difference between sequences to account for minor structural variation. Gene families were defined using a 50% identity threshold. Gene triplet forks (i.e., points in the pangenome graph where gene neighborhoods structurally vary between genomes) were also represented in binary format (`struct`).

*Proteins*. Annotated protein fasta files per isolate were clustered into panproteomes using CD-HIT^31^ (`v4.8.1`) with default parameters, except that description length limits were disabled (`-d 0`) to ensure that protein names were not shortened. Protein-level matrices were generated using both raw counts (`protein_count`) and binarized values (`protein_binary`).

*Domains*. Protein domain analysis was conducted using InterPro^32^ (`v5.74-105.0`), focusing on Pfam/Xfam^33^ annotations. Domains were similarly converted to count and binary matrices (`domain_count`, `domain_binary`).

All features were compressed as columnar sparse matrices in Parquet format to minimize file size, then linked through DuckDB views for efficient SQL querying of feature subsets. The compressed matrices were subsetted for each drug/drug class for subsequent model development.

### Machine learning models

To identify encoded features predictive of AMR phenotypes, we implemented a supervised ML framework using logistical regression (LR). In addition to their demonstrated ability to recover known AMR genes, LR classifiers^34^ were selected for their simplicity, interpretability, strong performance compared to baseline, and reasonable compute times. All models were built using the tidymodels ecosystem in R^34^ and applied across multiple molecular feature scales, species, and drugs/drug classes.

For each drug or drug class with sufficient data in each species, we subsetted the feature spaces to those genomes tested against that drug using DuckDB. We applied 5-fold cross-validation to each dataset with stratified sampling to preserve resistant/susceptible label ratios. We ensured that each fold had at least one observation of each label and enforced a minimum of 40 samples per model.

*Holdouts.* The `gene_count` matrices are stratified based on the country and year to develop separate models (e.g., *S. aureus* aminoglycoside model trained on US isolates, or a *S. aureus* methicillin model trained between 2010–2014). The holdout models were trained and tested on the self (e.g., US-trained, US-tested models) as well as on other held-out sets (e.g., US-trained, Thailand-tested; 2000–2004 trained, 2015–2019 tested models for a particular bug-drug combination).

#### Logistic regression

LR models were trained using glmnet^35^, with hyperparameters tuned over a predefined grid of penalty (λ) and mixing (α) values. The `mixture` hyperparameter ranged from 0 (ridge regression, L2) to 1 (least absolute shrinkage and selection operator, lasso, L1), with intermediate values representing elastic net models in increments of 0.2. Models were trained using a grid search over this hyperparameter space, and the best model was selected based on the Matthews correlation coefficient (MCC). A separate iteration of models underwent dimensionality reduction by principal component analysis (PCA) pre-processing, reducing features to principal components and retaining 95% of variance in the dataset.

Final performance metrics, particularly normalized MCCs (nMCC), were computed on held-out testing sets for comparison across feature types, drugs, and drug classes. The MCC accounts for all four elements of a binary confusion matrix: true/false positives and negatives in a single metric, which we have normalized to a 0–1 range for ease of interpretation. In this framework, an **nMCC of 1.0** indicates perfect prediction, **0.5** reflects random performance, and **0.0** represents complete disagreement with the true labels. Together, these values provide a balanced, interpretable framework for assessing predictive performance across drugs, drug classes, and molecular feature scales.

Following model training and evaluation, we extracted feature importance scores^36^. To prioritize biologically informative predictors, we ranked features by their absolute variable importance.

For each drug- or drug class-specific model, we save performance metrics, and the model features along with their corresponding importance values and signs. The latter are interpreted as the strength of feature associations with resistance (if negative) or susceptibility (if positive).

#### Label shuffling

To test model robustness, we performed random shuffling of AMR phenotype labels and retrained the models. Their performances with and without label shuffling were compared to evaluate their ability to discern biological significance in the real data.

#### Fisher’s exact test baseline

As a baseline comparison against ML, we performed feature-wise two-sided Fisher’s exact tests on binary genomic matrices representing feature presence or absence to identify individual molecular features associated with AMR. We selected a two-sided alternative hypothesis to capture both positively and negatively associated features. To account for multiple hypothesis testing, we applied Benjamini-Hochberg (BH) correction to control the false discovery rate (FDR). Adjusted *p*-values and binary significance calls (based on an FDR threshold, default *`Q`*`=0.05`) are reported in the *Results* section.

#### Annotations of the top features

To identify known AMR-associated features, top features were mapped to NCBI’s Reference hidden Markov model (HMM) Catalog of ARGs^37^ using HMMER `v3.4`^38^. While domains are species-agnostic, gene and protein annotations are not. To interpret these varied features across ESKAPE, genes/proteins/domains were also mapped to HMMs for clusters of orthologous genes (COGs^39^), enabling cross-species comparisons.

#### Cross-drug and MDR analyses

Cross-drug evaluation was done through pairwise comparisons across gene-, protein-, and domain-matrices such that each model was trained on data from one drug and tested on another (from the same or a different class). The datasets used for training and testing were mutually exclusive in the case of cross-drug testing.

Multidrug resistance (MDR) is defined here as resistance to more than one drug class and models to predict MDR have been developed following two distinct approaches; 1) *Multiclass classification*: MDR was considered a multiclass problem where each class represents a given combinatorial resistance profile—susceptible, resistance to single drug class (e.g., penicillins, fluoroquinolones), resistance to different combinations of multiple classes (e.g., lincosamides_macrolides_penicillins, cephalosporins_fluoroquinolones) as observed in each species. A multinomial logistic regression model was trained and tested for this framework. 2) *Multilabel classification*: MDR was treated as a multilabel problem, and each drug class was considered as an individual label. The multilabel classification was implemented using the binary relevance technique, where each label was treated as an independent binary classification task. This was performed using the glmnet algorithm through the mlr package^40^.

### Dashboard

We developed an interactive dashboard (https://jravilab.org/amr) to support exploration of AMR patterns by integrating manifold metadata with ML outputs. The dashboard is organized into four main tabs: `Metadata`, `Model Performance`, `Bug/Drug Feature Comparison`, and `Model Holdouts`. Each tab is designed to enable dynamic exploration of the data and results.

The `Metadata` tab of the dashboard enables users to visualize AMR surveillance data per bug by phenotype, country of isolation, host, and isolation source. These visualizations reveal both the diversity of resistance data and the heterogeneity of sampling contexts. As such, they suggest that resistance trends must be interpreted within a broader ecological and geographical framework. Complementing the plots, data cards display key summaries of genomes, drugs, and other relevant metadata.

The `Model Performance` tab allows users to explore the results of our pre-computed ML models and their predicted resistance phenotypes for specific drugs across different bacterial species. Users can customize their view by selecting a drug class, which filters the list of available drugs. After choosing a drug from the list and specifying a data type (binary or count), the dashboard displays a side-by-side comparison of model performance for each species. Boxplots show the distribution of performance scores for all models in the selected drug class, while overlaid points represent the performance of the selected drug’s models. This visualization allows users to assess how well models for a specific drug perform relative to the broader range of models in its class, using the model performance metric (e.g., nMCC).

The `Bug/Drug Feature Comparison` tab has two distinct options to visualize and compare important predictive features: across species and across drugs/drug classes. The models can be filtered based on molecular scale and data type for comparison. The “Across bug” option shows the ranked list of COGs representing important features from specific drug/drug class models in selected species, along with annotations and links to COG IDs (https://www.ncbi.nlm.nih.gov/research/cog/cog/) and NCBI HMM accessions (www.ncbi.nlm.nih.gov/genome/annotation_prok/evidence). The “Across drug” option also shows within-species features from selected drug/drug class models. The number of features to visualize from each model can be adjusted using a slider (up to 100).

The final tab, `Model Holdouts`, allows users to explore the performance and features of country and year holdout models tested on self and other datasets within a species and drug/drug class combination. This section provides insights into the exclusive and shared features across geographical regions and over time. The dashboard serves not only as a passive visualization tool, but also as a starting point for further investigation, guiding users toward the most informative genomic markers and robust prediction targets across molecular scales and data types.

## Results

### Compiling a comprehensive (meta)dataset

Accurate prediction of AMR from genomic data is a critical step toward enabling faster diagnostics and managing the spread of resistant infections. Public repositories like BV-BRC contain a wealth of bacterial genomes paired with AMR phenotypes, but variations in data quality and consistency across species and drugs pose challenges for systematic ML analyses. To address this, we curated a clean, multiscale dataset spanning ∼13,000 genomes with lab-confirmed AMR phenotypes across the ESKAPE pathogens, a group of clinically significant bacteria notorious for their multidrug resistance and global health impact. Genomic data and metadata were acquired for *Enterococcus faecium* (`n=1106`), *Staphylococcus aureus* (`n=3612`), *Klebsiella pneumoniae* (`n=5120`), *Acinetobacter baumannii* (`n=1710`), *Pseudomonas aeruginosa* (`n=1323`), and *Enterobacter* species (`n=499`). Constructing this dataset presented both an opportunity and a challenge: although all data were sourced from BV-BRC, assembling an ML-ready framework required extensive harmonization of genomic sequences, antimicrobial susceptibility profiles, and molecular annotations (**Figure 1**). We extracted and integrated multiple layers of information, including sample metadata (e.g., collection year and geographic origin per isolate) and multiscale molecular features, such as gene, protein, domain, and structural-level count and binary representations.

In total, our dataset spans 57 drugs (single or in combination) and 25 drug classes across the ESKAPE pathogens. The AMR data and associated metadata can be accessed via our dashboard (jravilab.org/amr, `‘Metadata’` tab), summarized by resistance phenotypes, drug classes, geographic origins, host types, and clinical specimen sources. Resistance profiles varied substantially, revealing both pathogen-specific biology and selective antibiotic pressures due to clinical history. For instance, *E. faecium* shows a high proportion of resistance to aminoglycosides and reflects how enterococci are relatively impermeable to these agents^41^. *A. baumannii* presents the most frequent overall drug class resistance proportion, particularly against fluoroquinolones, aminoglycosides, and cephalosporins (**Figure 2A**). Notably, resistance to last-line antibiotics like meropenem (475 resistant vs. 248 susceptible) was widespread and consistent with *A. baumannii*’s classification as a WHO critical priority pathogen (**Figure S1**). Resistance to older antibiotics like chloramphenicol and streptomycin was also observed, which indicates how multidrug resistance has become fixed in the population even after discontinuation of routine clinical use of these older drugs (**Figure 2B**). In contrast, resistance data for *S. aureus* were skewed toward the susceptible phenotype in recent years, which predominated in 12 of the 14 drug classes represented. This highlights *S. aureus* as an important counterpoint, where susceptibility remains more common, yet its clinical significance and potential for rapid resistance acquisition make continued surveillance essential. When looking at host source and geography metadata (**Figure 2C**; jravilab.org/amr, `‘Metadata’` tab), *A. baumannii* were predominantly sourced from human hosts, with the majority originating from blood, respiratory, wound, and tissue samples, from the US (`n=1107`), and other samples from Tanzania, Thailand, China, and Germany. Meanwhile, the *S. aureus* dataset was concentrated with isolates from the US and UK (`n=642`, `1115`, respectively), with less representation from Asia, Europe, and South America. The other ESKAPE pathogens were spread across the globe as well. These patterns likely reflect sampling emphasis and sequencing capacity rather than the true geographic burden of clinical AMR but nonetheless capture the breadth of ESKAPE resistance across hosts, geographies, and drug classes.

**Figure 2.**
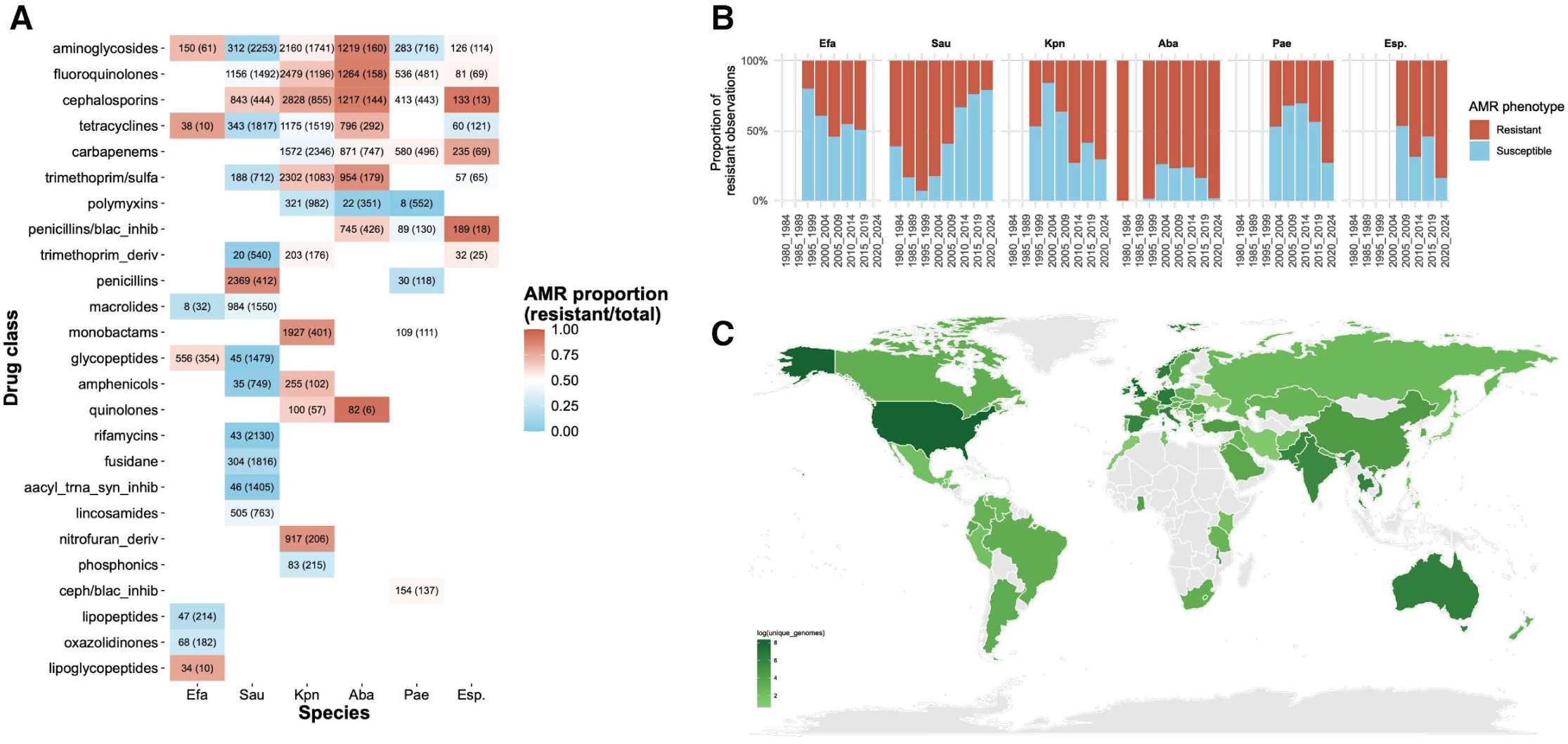
AMR prevalence across ESKAPE. **A.** The heatmap shows the fraction of genomes per species that are resistant to a particular drug class; red tiles indicate a higher frequency of reported resistance, while blue tiles indicate a greater fraction of susceptible genomes. Each tile shows the number of resistant and susceptible genomes (the latter in parentheses) for a given drug class with available whole genome sequences from BV-BRC. Rows are sorted by drug class data availability across species and the total number of observations. Only combinations with sufficient samples and class balance for our initial ML filtering are reported here (e.g., species-drug class combinations with fewer than 40 samples may be available through BV-BRC, but they are not shown). The drug class names are shortened for aesthetics; the complete names are listed in **Table S7**. **B.** The bar chart shows the proportion of resistant and susceptible isolates in each species across years, grouped into 5-year bins from 1980 to 2024. **C:** The choropleth shows the global distribution of unique resistant isolate counts across ESKAPE (log-transformed), where darker colors represent more genomes and lighter colors indicate fewer genomes.

This structured metadata landscape reveals distinct, lineage- and region-specific resistance trends across pathogens and highlights critical reservoirs of resistance both within and beyond clinical settings. It also supports targeted modeling efforts and enhances our understanding of AMR dissemination dynamics across ecological and geographic boundaries.

### Interrogating model performance across molecular scales and data types

Biological systems are organized across multiple molecular scales, from genes that encode genomic information and reflect heritable variation, to proteins that execute cellular functions, and to protein domains that represent conserved structural and functional subunits. Incorporating these three molecular scales into ML models allows for a more holistic representation of biological processes, enabling the detection of both broad genetic patterns and fine-grained functional features that may otherwise remain hidden if only a single scale is considered. Moreover, encoding these features as binary or count data types allows us to emphasize the presence or absence of key resistance-associated features and preserve the quantitative variation of these features, thereby enriching the information available to the models. Leveraging the value of this approach, we trained thousands of ML models to capture the essence of AMR prediction in 57 drug models and 26 drug class models spanning the ESKAPE species, three molecular scales (gene, domain, protein), and two feature encodings (feature counts vs. binary) (**Table S1**); each model yields ranked AMR feature lists per bug-drug (pathogen-antibiotic/class combination), providing a vast and comprehensive prediction framework that is easily accessible for exploration in our dashboard (jravilab.org/amr). The class balance of AMR phenotypes against individual drugs in the original BV-BRC dataset ranged broadly from 1% of *P. aeruginosa* genomes being colistin-resistant to 96% of *A. baumannii* genomes being cefotaxime-resistant (**Figure S1**). Such extreme examples with limited training data for ML were filtered out to prevent unstable predictions and poor generalizations. As a performance baseline, we generated randomized models by shuffling resistant/susceptible labels, severing the association between feature composition and biological outcomes. To account for variable class balance, we reported nMCC^42^ values to evaluate ML models, which encompass true/false positives and negatives in a single metric. The shuffled label models were expected to yield an nMCC=0.50 corresponding to random performance, and the observed median value of 0.49 confirmed this expectation. However, models trained on data with extreme class imbalance or small sample sizes exhibited noisier run-to-run variation in performance, which indicates that species-drug combinations with skewed class distributions should be treated with caution (**Table S2**). Across species, scales, data types, drugs and drug classes, most models showed excellent predictive performance (median `nMCC=0.89`), far surpassing random classifiers (`median nMCC=0.49`) (**Figure 3A**; jravilab.org/amr, `‘Model performance’` tab; **Figure S2; Table S2**), lending confidence to our models’ ability to uncover biological significance from multiscale data paired with AMR phenotypes. Interestingly, no single molecular scale yielded superior performance across the board, reflecting the biological contribution of each scale to AMR (**Figure 3B**). *E. faecium* showed uniformly high performance regardless of molecular feature scale, while in *S. aureus*, gene-level models performed better, albeit marginally, with shorter distribution tails, suggesting more consistent performance across models. *K. pneumoniae* and *A. baumannii* showed a range of moderate to high performance across drugs, scales, and data types. *P. aeruginosa* did the worst on all scales except `struct` (using genomic structure and proximal genes as their features). Finally, in *Enterobacter* spp. fewer high-performing models were found, which might be attributed to smaller datasets with a mixture of multiple species (**Figure S2**). Across ESKAPE and across drugs, nMCCs differed only slightly between binary (`median nMCC=0.89`) and count (`median nMCC=0.90`) data types, suggesting that the choice of data type has a limited impact on predictive performance (**Figure 3B**). In *E. faecium*, *S. aureus*, *K. pneumoniae*, and *A. baumannii*, both data types consistently yielded high nMCC values (typically `>0.90`), indicating that both binary and count-based feature encodings are viable for AMR prediction.

**Figure 3.**
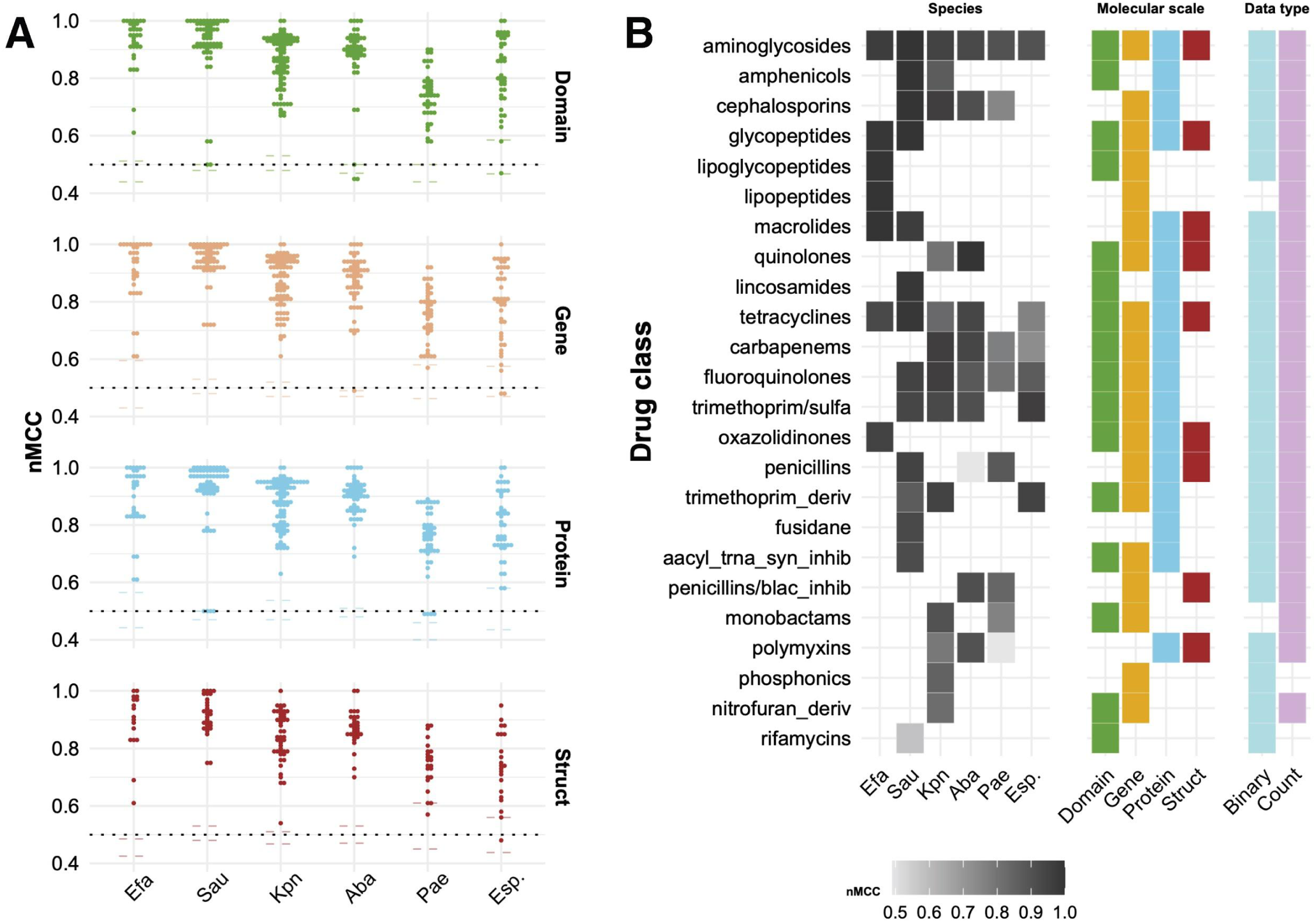
Model performances across ESKAPE. **A. Model performances against label-shuffled baselines.** Points in the plot represent the performance range of all drug and drug class models based on different feature types of a molecular scale (gene/protein/domain/struct) for one species. The dashed lines for each species and scale represent the median and interquartile range nMCC values for models where the AMR phenotype labels were randomly shuffled as a control to eliminate detectable biological signals between features and labels. These label-shuffled models represent random performance (nMCC ∼ 0.5; overlapping with the dotted 0.5 nMCC random performance line) and emphasize the prediction capabilities of the true models with biologically relevant labels. **B. Best performing feature scales across tested drug classes per species.** Grayscale colored tiles under ‘Species’ represent nMCCs for models, where black (1.0) shows perfect predictive performance, and light grey (0.5) indicates random performance. The drug classes are sorted based on their performance, and data availability across species. The next panels show tiles where that molecular scale or data type produced models with the top-ranked nMCCs. In cases where multiple molecular scales or data types are shown for a given drug class model, this indicates a tie in nMCC ranking. The drug class names are shortened for aesthetics; the complete names are listed in **Table S7**.

Despite having considerably fewer samples than features in our datasets—predisposing ML modeling to sparse matrices, overfitting, poor performance, and lack of generalizability—we found that dimensionality reduction models slightly underperformed compared to those using the full feature space (**Table S3**), and did so at a moderately increased computational cost (`∼20%` greater mean run times). Due to this and the desire to keep features interpretable, we did not pursue PCA models further.

### Predicting AMR in *Staphylococcus aureus* and *Acinetobacter baumannii,* a case study

We trained per-drug and per-drug-class models using both raw counts and binarized representations of gene, protein, and domain-level features within each bacterial species across the ESKAPE pathogens. Given the sheer breadth of the results, we use a focused case study of the evolutionarily distinct opportunistic pathogens *S. aureus* and *A. baumannii*, which differ in pathogenesis and physiology, to illustrate the kinds of insights our pipeline can offer. Both *S. aureus* and *A. baumannii* had ample data for ∼20 antibiotics each and models across gene, protein, and domain scales achieved consistently high performance above 0.70 median nMCC across most drugs in both species (**Figure 4**; jravilab.org/amr, `‘Model performance’` tab). This case study illustrates how our models capture both (i) species-specific biology for the same antibiotic and (ii) robust performance on distinct drug-species challenges. When we examined drugs tested on both, such as gentamicin, the models successfully captured species-specific resistance biology: in *S. aureus*, gentamicin resistance was linked to canonical aminoglycoside determinants across all scales (domains: PF13523, acetyltransferase domain; genes: [acetyltransferase](https://www.uniprot.org/uniprotkb/O52241/entry), [aacA-aphD](https://www.uniprot.org/uniprotkb/P0A0C1/entry); proteins: [acetyltransferase](https://www.bv-brc.org/view/Feature/PATRIC.1413508.3.JIZU01000026.CDS.294.1769.rev), [AAC(6’)-Ib/AAC(6’)-II](https://www.bv-brc.org/view/Feature/PATRIC.1413508.3.JIZU01000026.CDS.294.1769.rev))^43^ whereas in *A. baumannii* the top predictors included *[aadB](https://www.uniprot.org/uniprotkb/E9LWD7/entry)* genes with [phage integrase domains](https://www.ebi.ac.uk/interpro/entry/pfam/PF13495/), highlighting a different genetic basis for resistance. We also mapped features to clusters of orthologous genes (COGs), which allows us to traverse biological scales and bacterial species from specific resistance genes to broader functional domains^44^. This mapping revealed both conserved and species-specific elements of resistance. For example, COG1961 (a site-specific D recombinase SpoIVCA/D invertase PinE) was important for gentamicin and tobramycin resistance in both organisms, but has not been recognized as an ARG in the literature. However, the same COG was also uniquely important for ciprofloxacin, levofloxacin, and tetracycline resistance in *S. aureus* and for trimethoprim/sulfamethoxazole resistance in *A. baumannii*. Thus, the models not only detect the “right” resistance features, but also adapt to species-specific resistance repertoires. By contrast, when focusing on distinct drugs per bug, the models still performed strongly; tetracycline resistance (median `nMCC=1.00`) predictors in *S. aureus* included *[tetK](https://www.bv-brc.org/view/Feature/PATRIC.1280.11187.FQPS01000018.CDS.3.1406.fwd)* and *[tetM](https://www.bv-brc.org/view/Feature/PATRIC.1280.11187.FQPS01000018.CDS.3.1406.fwd)* genes and proteins (**Figure 4C**), and a plasmid replication initiation factor domain (<u>PF02486</u>) and [replication initiation protein](https://www.bv-brc.org/view/Feature/PATRIC.1280.39459.1280.39459.con.0014.CDS.3486.4439.rev) that implicates small plasmids, known as the dominant staphylococcal tetracycline resistance mechanism^45^. In *A. baumannii*, moxifloxacin is one of the best models (**Figure 4B**), and the top predictors for that fluoroquinolone resistance included [plasmid replication proteins](https://www.ncbi.nlm.nih.gov/Structure/cdd/COG5527), [transposases](https://www.bv-brc.org/view/FeatureList/?eq(plfam_id,PLF_469_00000003)#view_tab=features), and [prophage-associated hypothetical proteins](https://www.bv-brc.org/view/Feature/PATRIC.470.8027.470.8027.con.0004.CDS.98485.98898.rev). These findings validate the ability of our models to recover known resistance determinants from genome-wide feature sets and highlight the importance of mobile genetic elements (MGEs) in shaping AMR across both species.

**Figure 4.**
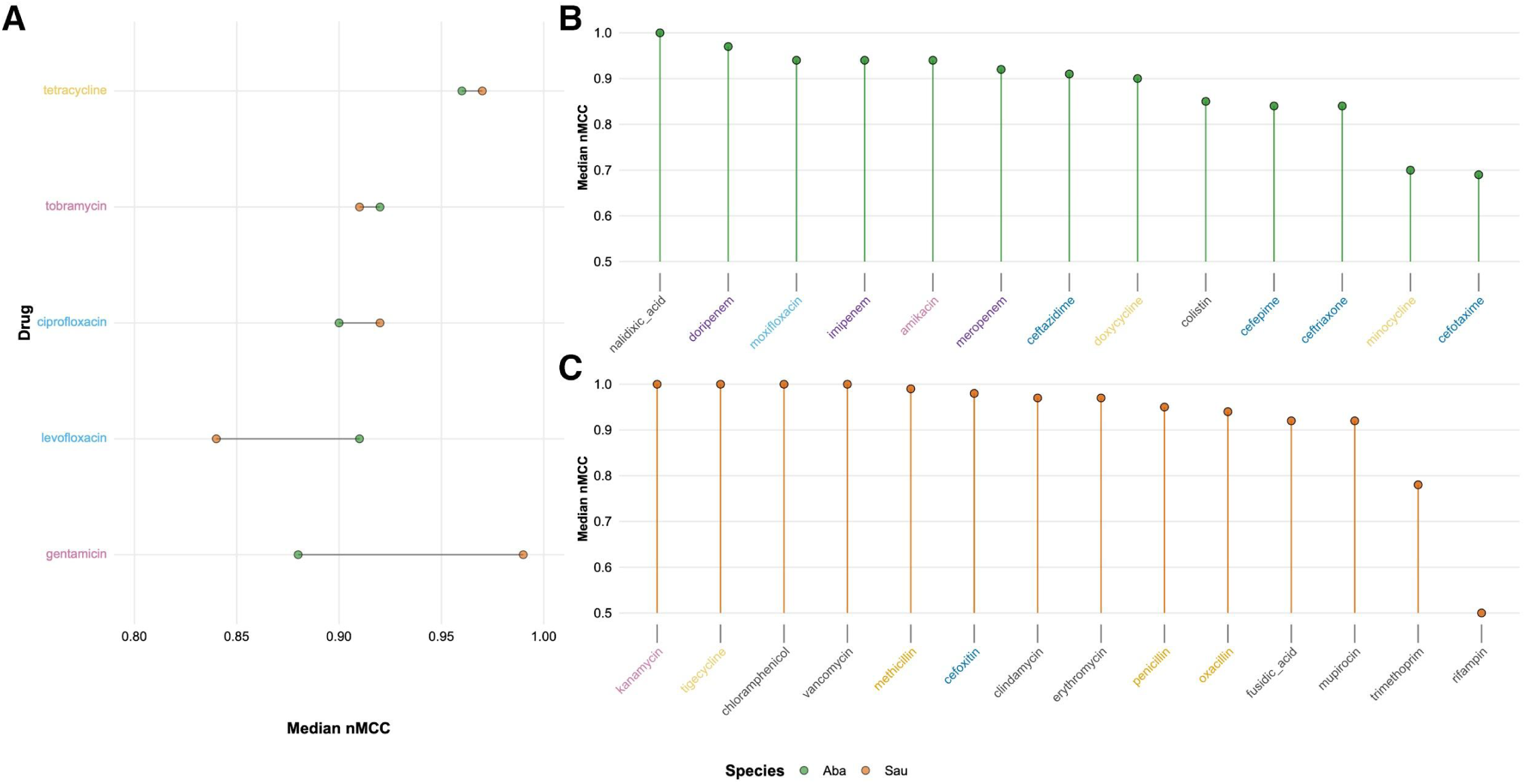
Comparison of model performance across *S. aureus* and *A. baumannii*. **A. Median performances of drugs used in both species.** Points indicate median nMCC values calculated considering all models specific to molecular scales and data types per drug for *S. aureus* (orange circles) and *A. baumannii* (green circles). The drugs are ordered based on the differences between the model performances. For drugs used in both species, performance metrics are comparable. **B & C. Per-drug model performance for antibiotics that are not shared between the two species**. The drug models in both species have shown robust performance, with the exception of cefotaxime and minocycline in *A. baumannii* **(B)**, while trimethoprim and rifampin in *S. aureus* **(C)**. The near-random median nMCC for minocycline resistance in *A. baumannii* is attributed to the domain-scale models (**Table S1**).

Known resistance determinants were consistently recovered–for example, [ErmC](https://www.bv-brc.org/view/Feature/PATRIC.1280.53227.CP150651.CDS.37731.38465.rev), [MsrA](https://www.uniprot.org/uniprotkb/P23212/entry), *[ermA](https://www.uniprot.org/uniprotkb/P0A0H3/entry)*, and *[ant(9)-I](https://www.uniprot.org/uniprotkb/Q7DIC3/entry)* for erythromycin resistance in *S. aureus*; or TetB, which contains PF07690, the largest secondary transporter superfamily, for minocycline resistance in *A. baumannii*. However, we also observed less-expected predictors. Many highly weighted features lacked functional annotation, such as “hypothetical proteins” or “domain of unknown function,” yet were frequently adjacent to known ARGs or MGEs. In *A. baumannii*, especially, top predictors included prophage-associated genes not described in CARD for these drug contexts. This was particularly striking for *A. baumannii*, where resistance profiles appeared to rely more heavily on poorly annotated genomic content, compared to the more known ARG-centric patterns in *S. aureus*. One possible explanation is that *S. aureus* — a historically well-studied and curated pathogen — has more complete annotation than *A. baumannii*, which may inflate the proportion of uncharacterized features in the latter. At the same time, the consistent genomic co-localization of these poorly annotated features with known ARGs and MGEs points to possible co-selection or regulatory linkage, suggesting they may simply reflect annotation artifacts but instead represent previously unappreciated phage- and MGE-linked pathways of resistance.

Some models yielded questionably perfect performance even with extremely imbalanced datasets – for example, vancomycin resistance was only observed in 3% of isolates, and subsequent models for *S. aureus* nearly all returned nMCC values of 1.00 (**Figure 4C**). While vancomycin resistance is sufficiently conferred by *van* operons, additional non-*van* features were assigned equal feature importance values as the resistance-determining features *vanA*/*B*; this result may arise from very small numbers of clonal resistant isolates yielding an array of MGEs (transposases, phage integrases, etc.) and genomic features that are potentially unrelated to vancomycin AMR (e.g., [thymidylate synthase](https://www.uniprot.org/uniprotkb/P0A0M5/entry), [dihydrofolate reductase](https://www.uniprot.org/uniprotkb/P0A017/entry), [DegV family protein](https://www.uniprot.org/uniprotkb/P13977/entry)) but are detected by models as associated. In other cases, reduced performance was expected: random baseline performance for rifampin in *S. aureus* is justifiable since the causative features of rifampin resistance are SNPs in the *rpoB* gene, and trimethoprim resistance arises from either dihydrofolate reductase mutations or acquisition of variant dihydrofolate reductase genes through horizontal gene transfer^46^. Collectively, these cautionary cases underscore that high model performance can be misleading under imbalance or SNP-driven phenotypes, and that meaningful biological interpretation requires considering both the predictive feature space and the underlying mechanisms before inferring generalizable AMR biology.

In *A. baumannii*, *S. aureus*, and our other analyzed pathogens, our approach yields many high-performing models backed by interpretable features across multiple scales that recapitulate known resistance mechanisms, emphasize the critical roles of MGEs in AMR, and expose underexplored features that may contribute to the adaptive landscape of AMR. These findings offer both mechanistic insight and avenues for hypothesis generation regarding novel resistance determinants. Performance results for all these models are available in **Table S1**, and both the performance metrics as well as contributing top features can be explored via our dashboard (jravilab.org/amr, `‘Model performance’` tab).

### Evaluating the robustness of our model performance

#### Testing model transferability across countries

Robustness is an essential, often overlooked, quality for ML models, and refers broadly to whether a model can still perform predictably while withstanding variations in the trends or quality of input data types. A model trained on historic samples from a certain population may perform exceptionally well when tested on samples from the same population, but it may overfit to only a narrow slice of biologically possible features. A robust model can be generalized to other settings and remains resilient to input data changes. For our AMR models to be useful for isolates from around the world and into the future, we asked whether and how features associated with resistance to the same drug or class vary with geographical location or temporal bins. Will our models be useful for isolates from newly sampled countries, or for samples collected five years from today? We trained models specific to countries and five-year bins to compare the context-specific models across drugs or classes. Further, to assess generalizability, we tested these stratified models on other datasets. As anticipated, the geographically stratified models for drugs tend to make better predictions on the same country test set than others, indicating the existence of regional AMR driving factors (**Figure 5A**). For example, we compare the country-specific models to predict lincosamide resistance in *S. aureus*. We report balanced accuracies (since MCC values were undefined when predictions were either all positives or all negatives) for the cross-testing performances across stratified models. In *S. aureus*, lincosamide data were available for four countries: Ireland, Singapore, Thailand, and the U.S.. The lincosamide gene model trained on *S. aureus* isolates from all countries taken together is a performant model with an nMCC and balanced accuracy (`bal_acc`) of 0.95. When the data were split by country (trained and tested on isolates from the same country), the performance increased for Ireland (`bal_acc=1.00)` and remained reasonably high for the U.S. and Thailand (`bal_acc = 0.90 and 0.83`, respectively); however, the model trained on Singapore isolates failed to predict susceptible samples (`bal_acc=0.50`). All country-specific models were further tested on isolates from the remaining held-out countries (e.g., **Figure 5C** reports these results for lincosamide resistance in *S. aureus*). Although all models performed poorly while testing on the U.S. dataset, the U.S.-trained model was successful in predicting resistance phenotypes across genomes from other countries, likely due to its well-connected stature and exposure to isolates from around the world and a significantly substantial sequencing capacity, allowing more diverse mechanisms of resistance to be captured in a larger dataset. The colistin model for *K. pneumoniae* performed modestly (`bal_acc=∼0.70`) when using merged data from 8 countries. In most cases, training and testing on single country data showed a performance deterioration over this value, but models limited to either Germany or Greece actually demonstrated a slight improvement. Regardless, for these and other single-country models, performance was not generalizable and performed worse when cross-tested, indicating the original combined models did identify signals that were lost when models were trained on single countries (**Figure S10A**). These data suggest that to discover robust regional patterns of AMR, we will require greater emphasis on sampling, sequencing, and data sharing through global public health collaborations.

**Figure 5.**
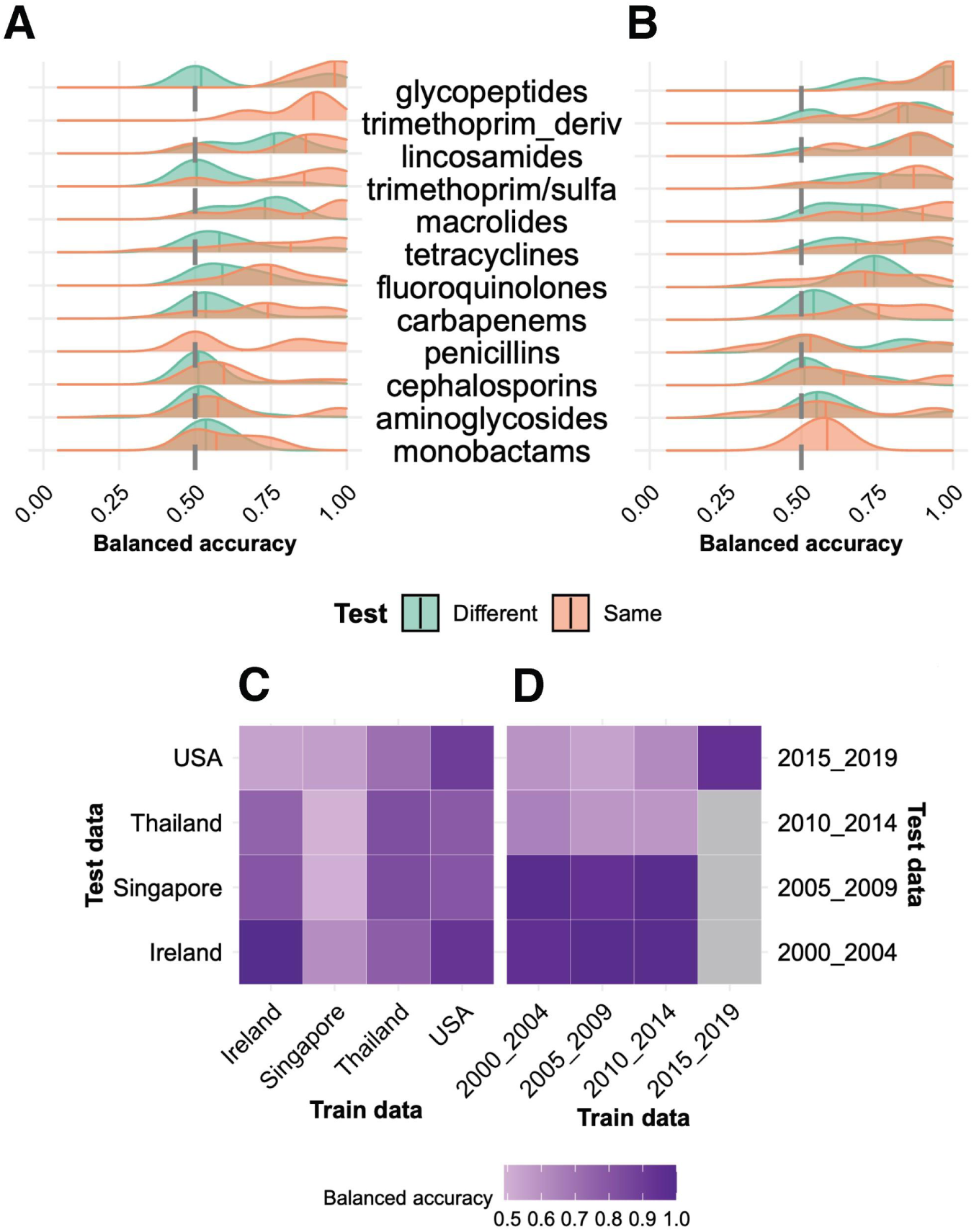
Geographic and temporal holdout performances. **A. The density of balanced accuracy from geographic holdout models.** We separated our training and testing data based on their metadata for country of origin, trained models on one country, and tested them against data from their own respective countries and all other countries. The performance of the geographic holdout models was evaluated on the same geographical dataset and other datasets. The plot shows the distribution of balanced accuracies of models across drug classes stratified by country and tested on the same country (orange) and different countries (green). The medians for each distribution are shown as vertical lines. **B. The density of balanced accuracy from temporal holdouts.** The models were stratified based on the 5-year bins of collection year, and models were evaluated on the same or different time periods. The plot shows the models performed similarly when tested on either dataset. The drug classes with sufficient data for both geographical and temporal stratification are shown in panels **A** and **B**. The drug class names are shortened for aesthetics; the complete names are listed in **Table S7**. **C. The performances of geographical holdouts for *S. aureus* lincosamide drug class models.** Others have established that AMR mechanisms and prevalence meaningfully vary based on their geographic origins. To assess how our predictive performance is influenced by country of origin, we demonstrate an example in *S. aureus* and the lincosamide drug class. **D. Temporal holdout testing for aminoglycoside drug class models in *S. aureus*.** As in A, we sought to discern the effects of sampling date on predictive performance and performed metadata-based holdouts of training and testing data. To demonstrate, we created four bins of 5-year intervals from 2000 to 2020, trained models on single bins, and tested them against other bins.

#### Testing temporal generalizations

Similarly, temporal holdouts and cross-testing on held-out time periods were performed. The collection years of the genomes were binned into 5-year intervals, and each dataset specific to a combination of species and drug or drug class was stratified into these bins. The general trend was that temporal-stratified models performed better when tested on the same time interval, with the exception of fluoroquinolones and trimethoprim derivatives, where cross-testing performances were equally good (**Figure 5B**). We demonstrate this trend with the aminoglycoside drug class in *S. aureus*, with AMR data from 2000 to 2019, resulting in four trained models (one per bin). Gentamicin was a common drug across these temporal bins; 2010–2014 and 2015–2019 groups had additional data for kanamycin and tobramycin, and 2015–2019 for streptomycin. The *S. aureus* aminoglycoside model was almost perfectly predictive with an `nMCC=0.99` and `bal_acc=1.00`; the temporally stratified models performed relatively well, too, with `bal_acc=∼0.94` (except for the 2010–2014 group, whose performance dropped to ∼0.60; see **Figure 5D**). This sudden drop in performance can be attributed to very low proportions of isolates with kanamycin and tobramycin resistance compared to gentamicin in the 2010–2014 data, leading to an unbalanced training/testing split. Each of the temporal models was further tested on other temporal groups to evaluate their robustness. Interestingly, all models performed poorly when tested on the most recent 2015–2019 dataset, likely due to shifting frequencies of AMR mechanisms. In the 2000–2004 model, aminoglycoside acetyltransferases—a known mechanism of aminoglycoside resistance—were two of hundreds of ranked features with approximately equal and low feature importance, indicating that while they were predictive, their contribution to AMR did not stand out in this time period. Conversely, the top two features in the 2010–2014 model were the same aminoglycoside acetyltransferases, but now in a much shorter list of top features and with ∼1.3–2.0x greater importance scores than the next two closest top-ranked features on the list, respectively, suggesting increasing prominence of these genes in AMR. Finally, in the 2015–2019 model, the same two features showed ∼8x greater feature importance than the next most important genes, reflecting their emerging dominant role in aminoglycoside resistance. These genes are prevalent in MGEs and plasmids that are often horizontally transferred. Another example is trimethoprim in *K. pneumoniae* with substantial data from 2005 to 2019. The stratified model for 2010–2014 (`bal_acc=1.00`) outperformed the overall trimethoprim model (`bal_acc=0.95`), whereas performances decreased for other holdout models (**Figure S10B**); interestingly, the 2010–2014 model also performed well on other temporally held-out datasets, and they did surprisingly better than their corresponding self-held-out. It is interesting to note that an integron integrase (*IntI1)* is the predicted top feature associated with trimethoprim resistance in *K. pneumoniae* overall, and this feature gains importance steadily from 2005–2009 to 2010–2014. This gene remains the top feature for the 2015–2019 model; however, other MGEs and genes associated with capsular polysaccharide synthesis join the top feature list. Their shifting importance over time reflects the establishment and fixation of this mechanism of resistance. The performances and top features of these and other geographical and temporal models are available in the dashboard (**Tables S4 and S5**; jravilab.org/amr).

#### Testing predictive transfer across drugs

Further, we investigated the generalizability of cross-drug models with our ML workflow. Training on one drug and testing on another drug examines how conserved resistance features are across drugs, both within and across drug classes. This approach highlights shared mechanisms (e.g., efflux pumps, modifying enzymes) that underpin multidrug resistance, while poor performance points to drug-specific determinants. We found that the molecular scale did not have a consistent impact on predictive performance (**Table S6**). Notably, models trained and tested on drugs within the same antibiotic class consistently outperformed those trained and tested on drugs from different classes across molecular scales and species (**Figure S3**). For example, in *K. pneumoniae*, models trained on cefotaxime performed better when tested on other cephalosporins than on other drug classes like fluoroquinolones. These findings support the need for class-aware modeling approaches and caution against applying models trained on one drug to predict resistance to mechanistically distinct drugs, except when identifying features involved in cross-resistance. In contrast, the oxacillin model in *S. aureus* performed exceptionally well while predicting resistance to drugs from other classes, including kanamycin and tobramycin (aminoglycosides) and clindamycin (macrolides) (**Table S6**). The top features shared among the oxacillin, aminoglycosides, and clindamycin models include glycerophosphoryl diester phosphodiesterase (ugpQ) and hydroxymethylglutaryl-CoA synthase (mvaS), which do not have a known relationship with AMR. However, they are often found within the *mecA* gene complex^47^, suggesting the association of *mecA* acquisition with multidrug resistance in *S. aureus*.

#### Establishing a statistical benchmark

To evaluate how well our ML models identify key resistance-associated genes, we compared the top-ranked features from each model to those identified by Fisher’s exact test, a standard statistical approach for detecting associations between gene presence and AMR phenotypes. Fisher’s test evaluates each feature in isolation, demanding a strong and consistent signal to withstand multiple testing corrections. Our ML pipeline, in contrast, evaluates features jointly: a predictor may not be strongly enriched on its own, but if it contributes unique information in the presence of other correlated variables, the model can assign it nonzero importance. In the gentamicin domain count models, nearly all the top-ranked predictors in each species were deemed significant by Fisher’s test, except in *E. faecium* (**Figure S4**). The overlap between Fisher’s and ML results confirms that both methods capture core resistance signals; yet, as expected, our ML pipeline additionally identified features such as the BRCA1 C-terminus domain (PF00533) and the phage replication protein CRI (PF05144) that appeared statistically insignificant in a univariate framework (**Figure S4**). Fisher’s test is too conservative for capturing the full spectrum of resistance-associated features, whereas our ML pipeline is better equipped to extract informative signals from correlated predictors. Together, these results demonstrate the necessity of incorporating ML to uncover nuanced, biologically relevant patterns that go beyond what is statistically enriched.

#### Assessing predictive resilience through feature elimination

As a final check for model robustness, we investigated model performance with feature subsets of decreasing size in our gene-level binary matrices. We started by training and testing with all genes for each species-drug combination as normal. Based on our ranked lists of top hits, we then removed those corresponding to the top 10% of all genes per matrix and retrained the models. We continued this process iteratively in increments of 10% (relative to the original total number of genes in each dataset) and tracked model performance until top predictive genes corresponding to as many as 80% of total genes had been removed. In cases where top hits comprised <10% of the total number of features in the dataset, genes were selected randomly without replacement until 10% was reached. We also fixed the LR regularization parameter to ridge regression; because lasso collapses clusters of genes with similar presence/absence patterns into singular representatives, it can lead to the repeated retention of the most resistance-associated genes across iterations of feature elimination. Such an approach results in the maintenance of high model performance even with near-complete feature removal, obscuring biological trends in our analysis (**Figure S5**). **Figure 6** shows nMCC vs. percent total features removed across ESKAPE for the top three antibiotics with 1) relatively balanced classes, and 2) a large number of observations (i.e., those with the greatest statistical power). Of note, we could not run this analysis on a few models due to large yet sparse feature matrices (e.g., many genes were present in only one genome). While nMCC decreased unanimously with increased omission of top genes as expected, we saw an intriguing sustenance of strong model performance with 10–20% removal in some instances;this is a non-trivial percentage that often corresponds to a few hundred genes. The lasso (**Figure S5**) and ridge (**Figure 6**) implementations of this feature elimination process both support a broad array of genomic modifications underpinning AMR, and not simply the presence of a singular AMR gene. While it is known that certain ARGs do unilaterally confer specific resistance profiles, a suite of adaptations to an AMR phenotype may accompany these ARGs, allowing the detection of the fingerprints of AMR even without the causative gene being sequenced.

**Figure 6.**
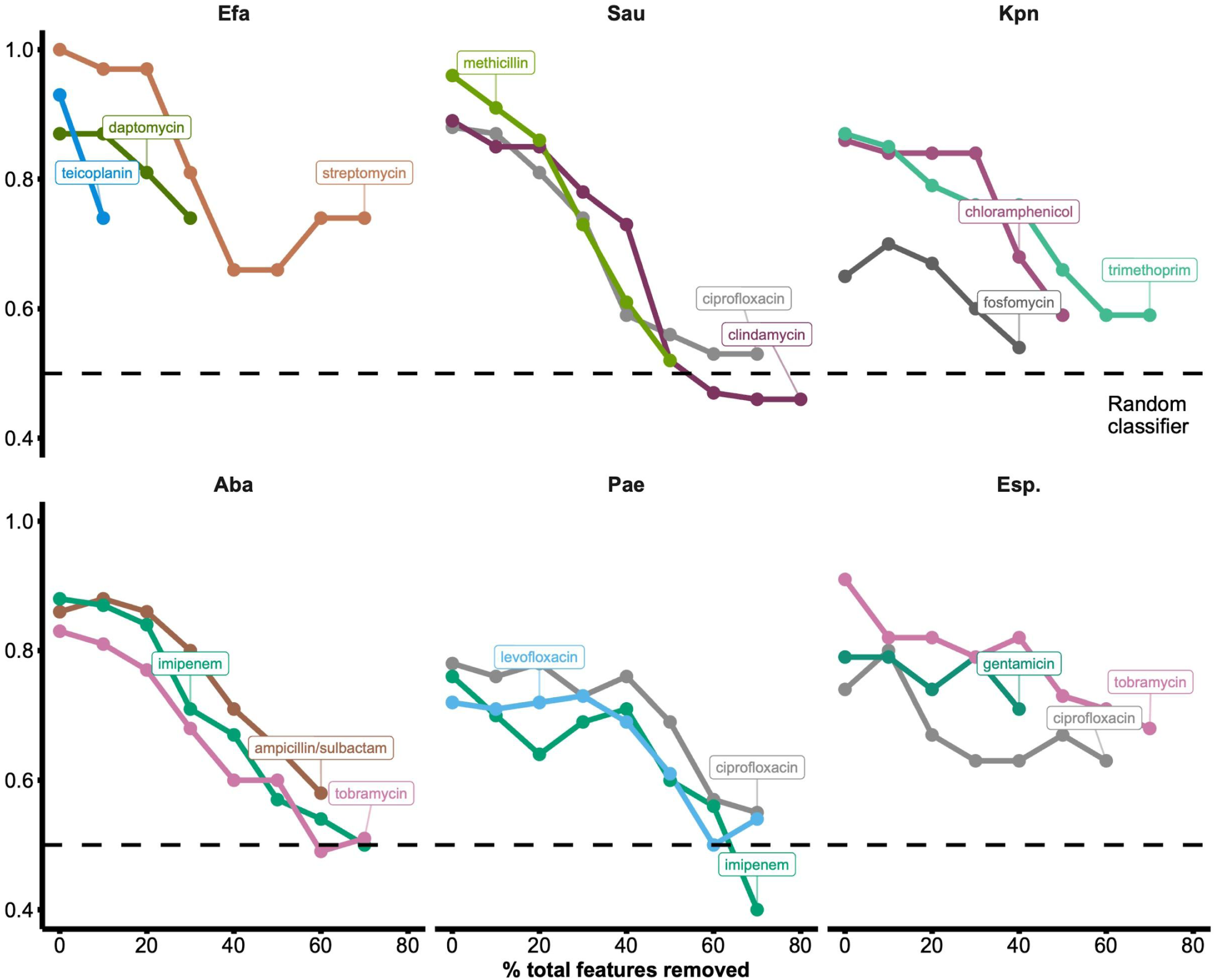
Removing top features iteratively to assess model robustness. To see how well models retain their predictive abilities when top features are removed, we started by training and testing with full datasets containing information about the presence/absence of genes. Once top features had been identified, we hid these from the models, retrained, retested, and repeated this process in increments corresponding to 10% of all genes in the respective datasets (relative to the original total number) per species-drug combination. In cases where all top features together comprised <10% of a dataset, genes were randomly selected without replacement for subsequent removal until 10% had been reached. Unlike with our other LR models, we fixed the regularization parameter here to ridge regression to prevent consolidation of our feature space to representatives from groups of genes with similar presence/absence patterns. For illustrative purposes, we show only the three drugs with the most observations in their respective smallest AMR phenotype classes per species. This reduces the noise in our results by filtering out drugs with high iteration-to-iteration variability. Performance, as measured by nMCC, falls uniformly with the removal of top features but often gradually, suggesting the presence of large sets of AMR-associated features.

### Discovering AMR mechanisms across species against a single drug class

We next asked whether the same drug selects for similar resistance mechanisms across phylogenetically diverse bacterial species. By comparing features of resistance, we uncovered shared evolutionary responses to antimicrobial pressure, which could inform surveillance strategies, predict resistance emergence, and guide cross-species stewardship efforts. The aminoglycosides class was uniquely suited for this analysis, since AMR data were available across ESKAPE (**Figure 7A**). We had isolates with AMR phenotypes for amikacin, gentamicin, kanamycin, streptomycin, and tobramycin (individual drug model performance shown in **Figure S6**). Using median nMCC, we compared all the drug models in the aminoglycosides class and the drug class model across ESKAPE pathogens. The aminoglycosides class performance in tested pathogens ranged from stellar for *S. aureus* (`nMCC=1.00`) to modest for *A. baumannii* (`nMCC=0.86`). All models showed strong performance with median nMCC > 0.80, with the exception of the *P. aeruginosa* and *Enterobacter spp.* amikacin models. In particular, models using domain or protein-level features generally achieved the highest median nMCC values. Gene-level models also performed well but showed slightly lower performance and greater variability across species (**Figure S7**). Structural variant features underperformed, which suggests that added feature complexity may not necessarily yield better predictions. Overall, species-level differences persisted, with *E. faecium, S. aureus,* and *K. pneumoniae* models mostly achieving high performance (`nMCC>0.90`), while *A. baumannii*, *P. aeruginosa,* and *Enterobacter spp.* showing more variability.

**Figure 7.**
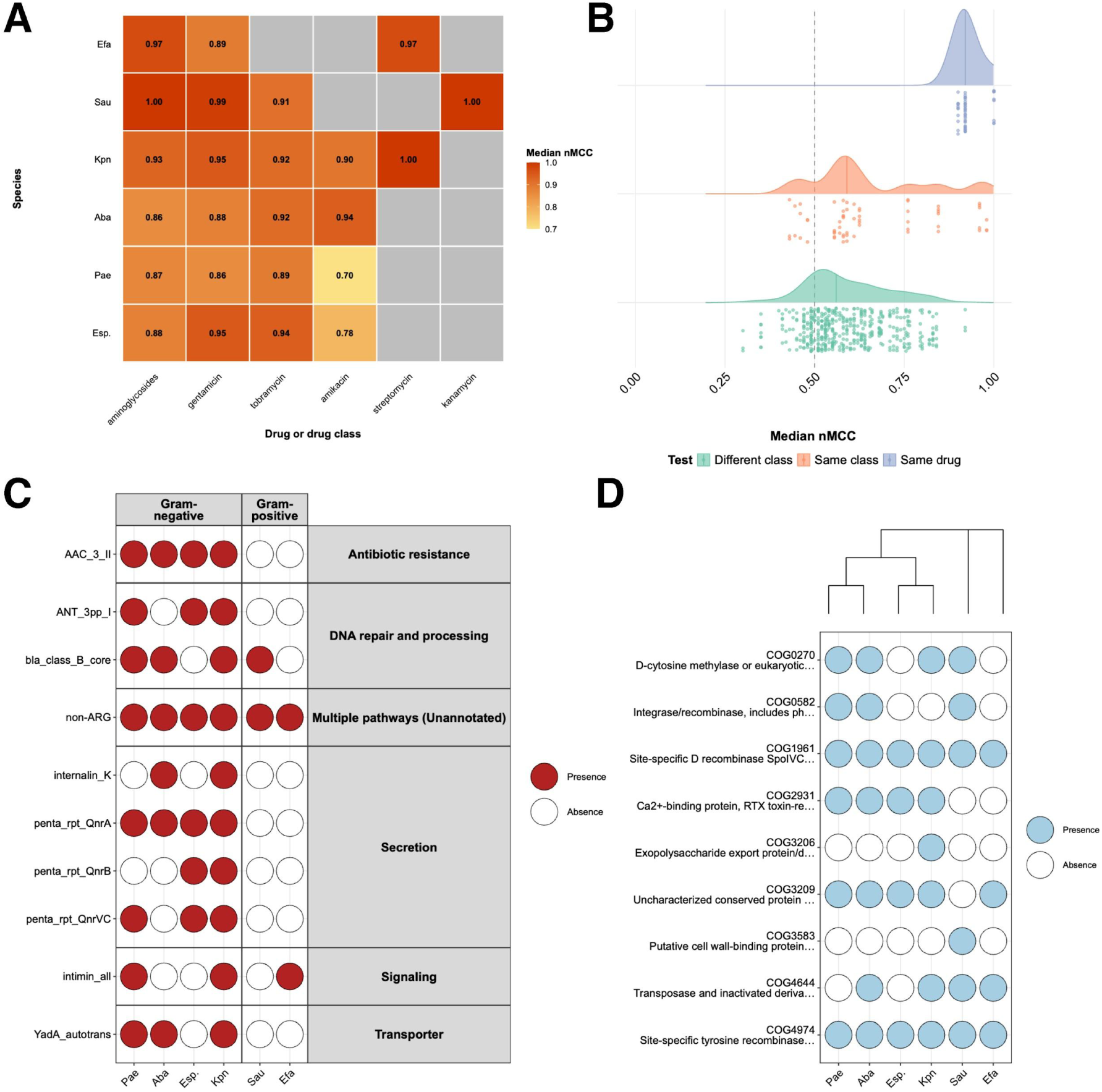
A case study of aminoglycoside models. **A. The median nMCC of aminoglycoside class and individual antibiotics from the class across ESKAPE.** The median nMCC was calculated for all models across molecular scales and data types per species. Higher performing models are shown in darker shades. Gray tiles indicate insufficient observations of that species/drug combination for modeling. **B. Cross-testing performances of aminoglycoside models.** The distribution of model performance across molecular scales for count data types are shown for aminoglycoside drug models tested on self (blue) vs. other drugs within (orange) or outside (green) the aminoglycoside class. **C & D.** We ranked all detected ARGs **(C)** and COGs **(D)** by total occurrence across all bug-drug-domain model combinations in the aminoglycoside class and plotted the top ten features. **C. Distribution of the most frequently occurring Antibiotic Resistance Genes (ARGs).** The ARG panel shows the presence (red) or absence (white) of ARG targets grouped by functional category (right) and split by Gram type of the bacterial species from the literature. Many ARGs are broadly conserved across Gram-negative species but largely absent in Gram-positives, though it is important to note that some features may be present across all species but may be ranked lower and therefore not plotted. **D. Distribution of the most frequently occurring Clusters of Orthologous Groups (COGs).** The COG panel shows the phylogenetic tree (generated by 16S analysis) of the six ESKAPE pathogens alongside the presence (blue) or absence (white) of the top COGs.

It is important to note that although nMCC in *S. aureus* for gentamicin was near perfect, such exceptional performance can be a warning sign of overfitting or extreme class imbalance. However, in *S. aureus*, these suspiciously high values are well-supported by high feature importance scores for top predictive features that are known causative mechanisms of aminoglycoside resistance (i.e., acetyltransferase domains, genes, and proteins). *E. faecium*, *P. aeruginosa*, and *A. baumannii* returned more moderate performance metrics for gentamicin, but also showed highly imbalanced resistance distributions (e.g., *A. baumannii* resistance proportion `≈ 0.87`) (**Figure 7A and S1**). These data are reminders that class imbalance can inflate performance metrics, and it is critical to consider resistance prevalence when interpreting model accuracy, and to pick the right evaluation metric based on what we are penalizing/optimizing for — such as the ability to accurately predict resistance of a new strain (to prescribe the best drug class). These findings underscore the need for balanced and sufficiently large datasets to ensure fair cross-species comparisons of predictive performance. The individual drug models, when evaluated against other drugs from the same class (aminoglycosides) and other classes, performed as expected; the model performances decreased when shifting from the same to a different class (**Figure 7B**). This observation supports that even though the aminoglycosides class-specific features like aminoglycoside acetyltransferases are identified among the top features for individual drugs, each drug model has identified the specific family of the enzyme that has better specificity for the drug, such as AAC(3)-1 that has better specificity for gentamicin, while AAC(6’)-I has better specificity for tobramycin^48^.

At the feature level, we mapped the top features to existing ARGs from the National Database of Antibiotic Resistant Organisms (**Figure 7C**). Among domain binary models for aminoglycosides, [ANT(9)](https://card.mcmaster.ca/ontology/39030) was identified as a top feature in *E. faecium*, *P. aeruginosa,* and *Enterobacter* spp., indicating shared resistance mechanisms against aminoglycosides. Other shared non-aminoglycosides ARG hits included *[mecI](https://card.mcmaster.ca/ontology/36263)*, the transcriptional repressor for *mecA* and cause of methicillin resistance, and [qac_SMR](https://card.mcmaster.ca/ontology/36004), the small MDR antibiotic efflux pump. While these are distinctly different mechanisms (e.g., specifically targeting methicillin vs. generic drug efflux-mediated resistance), they may serve as MDR markers that effectively predict gentamicin resistance as well. Among COGs (**Figure 7D**), we identified numerous phage, plasmid, transposase/insertion sequence elements across multiple species. These COGs have not been implicated in AMR directly, but most are linked to HGT, which underlies typical aminoglycoside resistance. We emphasize here an advantage our ML models provide over methods using only known ARGs: we can detect AMR acquisition mechanisms, adaptations, and MDR features (see dashboard; jravilab.org/amr); thus, helping expand our predictive potential and our biological understanding of resistance.

#### Fingerprinting AMR against multiple drug classes with MDR models

To address distinct aspects of MDR, we employed two modeling approaches (see *Methods*): i) *multiclass* classification, which assigns each possible combination of resistance and susceptibility across drug classes as distinct modeling classes to investigate how resistance co-occurrence (e.g., susceptible to all; resistant only to penicillins; resistant to fluoroquinolones_macrolides_penicillins; etc.); ii) the *multilabel* approach to independently predict resistance to each antibiotic class (e.g., penicillins, fluoroquinolones) within a single genome.

As a case study, the MDR multiclass model for *S. aureus* performed well with an nMCC of 0.87. Several of our *S. aureus* genomes reveal resistance to a single antibiotic class, including penicillins, tetracyclines, macrolides, fluoroquinolones, and cephalosporins, whereas the majority of genomes in *K. pneumoniae, A. baumannii*, *and Enterobacter spp.* exhibit resistance to two or more classes (**Figure S8A**). Interestingly, the multiclass model of *S. aureus* misclassified many of the susceptible genomes and multidrug-resistant genomes as penicillin-resistant (**Figure S8B**). This suggests a potential bias of overrepresentation of penicillin resistance features in *S. aureus,* which is often pointed out in the literature^49^. This also suggests evolution of MDR *S. aureus* strains often begins with penicillin resistance, which is also evident from the historical burden of methicillin-resistant *S. aureus* globally. Additionally, a small number of genomes labeled as susceptible to all antibiotic classes were incorrectly predicted as resistant, indicating the drawback of the antibiotic susceptibility testing, which does not include all possible drugs. Cross-class prediction errors were also observed, particularly involving fluoroquinolones—some resistant genomes were predicted as susceptible and vice versa. Fluoroquinolone resistance is typically driven by point mutations in genes such as *gyrA* and *gyrB*, which are not captured by gene presence/absence or abundance-based models. The multilabel model, which was built on genomes with AST data for all the tested antibiotic classes, reflected this limitation too, where fluoroquinolone resistance was not predicted at all, despite those genomes being resistant to multiple classes, including fluoroquinolones.

#### Questions enabled by our approaches

Our data dashboard aids navigation through these vast, curated AMR datasets across the ESKAPE pathogens. The dashboard includes different sections allowing users to explore AMR isolate data and our AMR prediction models. The `Metadata` tab illustrates the distribution of genomes across species, geographical location, and years. The different sections dedicated to prediction models allow visualization of performance and predictive features of multiple drug or drug class models in one or more species. The dashboard also includes geographical and temporal holdouts and cross-testing model performances and important variables. Our analysis, available in an interactive, easy-to-use manner via the dashboard (jravilab.org/amr), enables us to answer questions, such as: Which molecular scale performs better than others in each species-drug combination? What are the features conserved across molecular scales for a particular drug or drug class resistance? How do the features vary among drugs of the same drug class in a species? What are the shared or unique features across bacterial pathogens for the same drug or drug class? Is a model trained on a particular country or time point able to predict on data from other countries or newer time frames? Do the important features change over time or vary with geography?

## Discussion

Our investigation yields significant insights into the genetic underpinnings of AMR in the ESKAPE pathogens, demonstrating the predictive capabilities of ML across diverse bacterial species, drugs, drug classes, molecular scales, and data types. Models trained on molecular features at individual biological scales consistently achieved high predictive performance across pathogens.

### Consistent performance across most molecular feature scales

Across species and drugs, we observed roughly equivalent performance for different feature scales (**Figure 3B**). In contrast, the `struct` models tended to underperform slightly compared to other scales. They also required more compute time and yielded larger, less interpretable lists of predictive features. Domain models outperformed all models when it came to run time; they ran ∼50x faster than gene structure and ∼43x faster than the protein features from which the domains were originally extracted **(Figure S9**). This comparison does not factor in the extensive time to cluster the protein space through CD-HIT or to extract domains from InterPro, but it does illustrate the efficiency of domain-based models without sacrificing performance in the tested species. It is important to note that not all proteins have annotated domains, and if such proteins yielded resistance, they would escape detection by a domain-only approach. Additionally, binary and count models with features across molecular scales typically performed equivalently in the tested species, and when excluding the binary-only gene structure models to allow for a one-to-one comparison, the model runtimes across other scales were nearly identical. Given the potential that specific forms of AMR could be modulated by gene copy number, we may give an edge to count models for their ability to detect such differences, but they yielded few advantages in our testing.

### Top AMR features

We find that for most models, known ARGs constituted only a minor fraction of the top predictive features across our studied species, despite their undisputed clinical relevance. This challenges the conventional reliance on canonical ARG databases as the sole source for accurate resistance prediction. Instead, our models also find predictive value in other genomic elements described below, which could impact resistance through mechanisms such as gene regulation, epistasis, or co-selection. For example, regulatory changes can influence AMR by increasing the expression of efflux pumps, helping bacteria evade antibiotic effects even without changes to the underlying genes. Epistasis may affect AMR when a compensatory gain or loss of a gene or mutation restores bacterial fitness while preserving resistance. Co-selection occurs when genes linked to AMR are maintained in a population due to selective pressures from unrelated agents like metals, promoting resistance even in the absence of antibiotic exposure, as evidenced by the ubiquitous appearance of features involved in arsenic and mercury resistance across species and drugs.

Hypothetical proteins, genes, and domains of unknown function were common contributors to accurate resistance predictions, sometimes comprising over half of the top predictive genes. The prevalence of uncharacterized loci highlights a substantial knowledge gap in our understanding of AMR and presents a compelling opportunity for novel discovery in their AMR roles.

Features associated with HGT and MGEs were highly prevalent among predictive features as well. This aligns with the well-documented key role of MGEs in the acquisition and dissemination of resistance in taxa such as *S. aureus* and *A. baumannii*. When analyzing the top features driving AMR, we must consider the critical roles that gene flow within and across diverse taxonomic groups plays. While Pfam domains can be applied readily for cross-species analysis, comparing wildly variable gene or protein names can introduce problems even within a single species. To work around the roadblocks to understanding which features are common drivers of AMR across diverse pathogen species, we leveraged the power of HMMs for distant homology detection through HMMER and an HMM database of COGs^39^. By assigning gene or protein clusters to COGs, we bridge phylogenetic distance and find readily comparable features (**Figure 7D**) across scales.

Nonetheless, our models often successfully identified established AMR mechanisms (**Figure 7C**). A direct and compelling validation for known AMR mechanisms was observed in *S. aureus*, where we consistently identified acetyltransferases as the top two predictive features for gentamicin resistance across both gene and protein models (using counts or binarized features). Furthermore, the GNAT acetyltransferase^50^ domains contained within proteins were also the most predictive for domain count and binary models. As such, our models directly align with established mechanisms of aminoglycoside resistance, where aminoglycoside acetyltransferases are known to chemically modify the drug to prevent its ribosomal binding and confer resistance^50,51^. Our models consistently recapitulate known AMR mechanisms alongside novel ones.

Further analysis revealed patterns suggesting the prominence of HGT, a well-known method by which pathogens acquire AMR. Several high-frequency COGs, such as recombinases, occur in nearly all species, suggesting a strong role for mobile genetic elements, while some COGs show more restricted phylogenetic distributions, indicating possible lineage-specific adaptation (**Figure 7D**). These frequency-based rankings of ARGs and COGs highlight highly conserved resistance determinants and mobile genetic factors across ESKAPE. We iteratively removed top features from genomes and saw persistent predictive performance even with 20% or more of all features eliminated (**Figure 6**), leading us to suspect that HGT was playing a role. Pathogens share large genomic islands and other MGEs carrying ARGs, resulting in a high number of genes that are associated with but not causative of AMR. This phenomenon was likely reflected in a few species-drug combinations tested in our study, such as streptomycin in *E. faecium* or chloramphenicol in *K. pneumoniae*. For these, model performance did not drop until >20–30% of total features had been removed. An alternative explanation for this, besides HGT, is that selective pressures imparted by AMR acquisition and maintenance could predispose pathogens towards broader genomic and metabolic adaptations, leaving fingerprints of resistance well beyond the genetic feature(s) responsible for AMR. Meanwhile, performances for other species-drug combinations, such as teicoplanin in *E. faecium*, declined more rapidly, likely owing to small groups of genes (e.g., the *van* gene clusters) that directly confer resistance to glycopeptides. Once these are hidden, the remaining teicoplanin mechanisms and resistance signals that are SNP-based are difficult to discern from these models. Such resistance profiles tied to a handful of genes are easier to characterize in the lab but overall represent an exception rather than a rule in the world of AMR. This reiterates the need for ML given its advantage over traditional approaches in capturing the complex evolutionary shifts that occur in pathogens as they acquire resistance.

### Advances in the AMR field

ML approaches are rapidly growing as pivotal tools in addressing the critical global health threat of AMR by offering novel avenues for prediction and intervention^52^. These sophisticated models harness a diverse array of data from clinical patient information and detailed genomic sequences to microbiome insights and epidemiological trends^53,54^. Significant successes have been achieved by prior ML approaches in predicting AMR. Notably, ML algorithms such as XGBoost have predicted carbapenem-resistant *Klebsiella pneumoniae* infections in intensive care units with an average accuracy of 83%^55^. Other ML models have also shown competitive performance in predicting AMR in *Acinetobacter baumannii* directly from *k*-mer features extracted from WGS data, achieving essential agreement and category agreement exceeding 90% and 95%, respectively^56^.

Despite these compelling successes, prior ML approaches for AMR prediction face several significant limitations. A primary concern is data availability and specificity to particular pathogens and antibiotics. Many models are developed using exclusive data from a single hospital population, thereby lacking diverse, representative training data. More often than not, single-species models perform poorly even on closely related species or fail to generalize across different drug classes^57^. This necessitates rigorous external and prospective validation to ensure their adaptability and generalizability to other clinical settings. Moreover, current AMR prediction models are often constrained by their reliance on predefined resistance genes, limited feature types (e.g., only SNPs or ARGs), or narrow contexts, stalling the discovery of novel AMR mechanisms and posing risks of misclassification when using rule-based methods to assign AMR in resistant populations that do not carry the majority AMR genotype. Our approach addresses this by integrating multiscale molecular features derived from bacterial pangenomes, mapped to species-agnostic COGs and domains that aid in translating knowledge across species while capturing both canonical and noncanonical resistance signals. By moving beyond curated ARG databases and incorporating signals from mobile elements, hypothetical proteins, and structural genomic motifs, this work expands the feature space and allows the identification of emergent resistance determinants that would be missed by rule-based or single-scale methods. Our workflow is generalizable across available bacterial data and permits granular AMR insight and discovery; all our data, models, and results can be explored here: jravilab.org/amr.

### With a grain of salt

Although our models deliver strong, insightful results across pathogens, drugs, drug classes, and molecular scales, we must consider some inherent limitations. First, our metadata is dominated by high-resource countries, single-site or outbreak-focused studies, and human hosts, leaving a substantial portion of the AMR story untold (**Figure 2C**; jravilab.org/amr, `‘Metadata’` tab). **Figure 5A** demonstrates how models trained on isolates from one country and tested on isolates from other countries can yield dramatic performance shortfalls. These stark differences are essential to consider given the disparity in public data availability with broad geographic distribution: regions like South Asia, sub-Saharan Africa, and parts of Latin America exhibit some of the highest AMR-related mortality globally, as reported by a 2024 systematic analysis^4^, but often have no available genomic data with paired AMR phenotypes. A smaller but notable subset of isolates was sourced from animals, including pigs, dogs, and poultry. A One Health-aware surveillance framework that incorporates such data is necessary, especially considering the zoonotic potential of resistant *S. aureus*^58^ and *P. aeruginosa*^59,60^ strains. Further, given the even greater volume of antibiotics used in livestock than in humans, often for non-medical purposes^61^, future analysis of AMR patterns requires much more extensive sequencing and phenotyping of isolates from animal hosts.

Our molecular feature scales do not capture single nucleotide-level variations. Such variants are critical to certain types of AMR, and we must caution that our models are not designed to identify every possible form of AMR that may arise. However, even for drug resistance profiles that are dominated by SNPs such as in fluoroquinolone drugs (where resistance is typically caused by target site mutations^62^), we still observe some highly accurate models with predictive performance well above random baseline (**Figure S5**). To our initial surprise, when predicting fluoroquinolone resistance in *K. pneumoniae*, we saw an abundance of aminoglycoside-modifying enzymes (e.g., *aac6-IIc, aac6-Ib*), a typical cause of resistance against aminoglycoside drugs and a common feature observed in our models for those drugs. However, it has been shown that some aminoglycoside acetyltransferases have acquired mutations that allow them to act upon both aminoglycosides and fluoroquinolones like ciprofloxacin^63,64^ – this suggests that even without incorporating SNP-level information, we may still capture the effects of these mutations on higher-level phenotypes. Among top features, we also noted *bla* genes (encoding beta-lactamases) that often propagate with quinolone resistance genes (*qnr*) through mobile genetic elements^63^, suggesting that some top features are effective proxies for specific types of resistance even if they are not directly responsible for the effect.

Scarcity of phenotypic data and severe class imbalances for select antibiotics (and classes) in some bacterial backgrounds also complicated model predictions (**Figure S6**). In such cases, performance fluctuated between unreasonably strong (`nMCC=1`) and random baseline performance (`nMCC=0.5`), with many combinations lacking sufficient data for ML modeling in the first place. Many of the antibiotics classified under the WHO AWaRe framework (Access, Watch, and Reserve) are represented in our dataset. However, newer and last-line antibiotics remain underrepresented, limiting the availability of sufficient data for robust ML analyses. Cross-species consideration is important but was challenging in our analyses due to inconsistent cluster annotations between pangenomes, especially for gene and protein feature scales. As a workaround, we added high-level COG annotations for cross-species comparisons. Given the scarcity of a hypothetical pan-species pan-feature matrix and the computational burden that would be involved in constructing it, innovative methods are needed to bridge the cross-species gap in the future. The vast space of hypothetical proteins with no functional assignment represented an additional limitation for the depth and breadth of our analyses. Finally, future methods are warranted for training models to make predictions on newly sequenced genomes without the need to generate entire pan-feature matrices afresh.

### Impacts

All considered, this work demonstrates the utility of interpretable, multiscale ML models in uncovering novel resistance-associated features. It opens the door to mechanistically informed AMR prediction, where domain and protein-level insights point toward candidate resistance genes and genomic contexts. This framework supports a new paradigm for AMR research: one in which ML is not merely a black box classifier but a hypothesis-generating engine that directs experimental validation and surveillance strategies. We bolster this approach through our interactive data dashboard, allowing others to accessibly browse and interpret data from hundreds of models and thousands of isolates. In the future, we plan to integrate transcriptomic or proteomic data, when available, to refine predictions and use transfer learning across species to identify conserved or convergent resistance mechanisms. Transcriptomic or proteomic data could be incorporated to capture condition-specific resistance responses and post-transcriptional regulation, which are often missed by genome-only approaches. Furthermore, the use of transfer learning may reveal conserved or convergent resistance mechanisms across phylogenetic distances. This is especially relevant for identifying pan-species biomarkers of resistance and for preempting the spread of resistance traits in newly emerging threats. Finally, coupling this computational pipeline with real-time genomic surveillance efforts such as those run by public health laboratories could provide actionable insights into emerging resistance patterns. As more high-resolution phenotype-genotype datasets become available, we anticipate a virtuous cycle where ML-derived hypotheses drive experimental discovery, which in turn, enhances the predictive power and interpretability of future models.

## Conclusion

Our multiscale machine learning framework offers a powerful and interpretable solution to one of the most urgent challenges in modern medicine: the rapid and reliable prediction of AMR. Using publicly available data to construct pangenomes for various bacterial pathogens and applying established ML methods, we present a robust set of models that, using only resistant/susceptible labels for individual genomes, predict genes associated with drug or drug class resistance *de novo*. We share our data, metadata, and results through an interactive data dashboard available at jravilab.org/amr. nMCC values support the use of these models in correctly predicting susceptibility to different antibiotic classes from complete genome and WGS data. Many of our top feature hits have plentiful literature support (e.g., *S. aureus mecA* for methicillin resistance, *ermC* for macrolide resistance), while others are associated with pathogenicity islands and other mobile genetic elements that include resistance mechanisms (e.g., *S. aureus ugpQ* in the SCC*mec* locus). We also identify many features that are potentially novel AMR contributors. As genomic data continues to expand, our framework serves as a timely and adaptable tool for improving AMR surveillance, guiding therapeutic decisions, and accelerating the development of new diagnostics and interventions.

## Acknowledgments

We would like to thank Drs. Arjun Krishnan, Chris Mancuso, Emily Meyer, Jerome McKay, and Inka Sastalla for their invaluable feedback and support throughout this project. We would also like to thank Katerina Terwelp for reviewing our manuscript and its accompanying dashboard during development. Our project relied on the numerous resources shared through BV-BRC’s database and command line interface tools. This work utilized the Alpine high-performance computing resource at the University of Colorado Boulder. Alpine is jointly funded by the University of Colorado Boulder, the University of Colorado Anschutz, Colorado State University, and the National Science Foundation (award 2201538).

