## Supplemental Figures and Legends for "From sequence to signature: Machine learning uncovers multiscale feature landscapes that predict AMR across ESKAPE pathogens"

**Supplementary figures** 2

Figure S1. AMR prevalence across pathogens by drug class 2

Figure S2. Best performing feature scales across tested drugs per species. 3

Figure S3. Cross-drug model performance across species, molecular scales, and drug classes. 4

Figure S4. Demonstration of top ML features against Fisher's features. 5

Figure S5. Removing top features iteratively with lasso regression. 6

Figure S6. Best performing feature scales across tested drugs per species. 7

Figure S7. Aminoglycosides model performances across molecular scales for ESKAPE. 8

Figure S8. Proportions of predicted resistant classes across drug combinations. 9

Figure S9. Model runtime comparisons across feature scales. 10

Figure S10. Geographical and temporal holdout testing performances. 11

**Supplementary tables** 12

Table S1. Model performances across ESKAPE species, molecular scales, data types, drugs, and drug classes. 12

Table S2. Performances for shuffled baseline models. 12

Table S3. Performances for PCA models. 12

Table S4. Performances for geographical holdout models. 12

Table S5. Performances for temporal holdout models. 12

Table S6. Performances for cross-drug trained models. 12

Table S7. Shortened drug class names. 12

Table S8. Abbreviated drug class names. 12

#### Supplementary figures

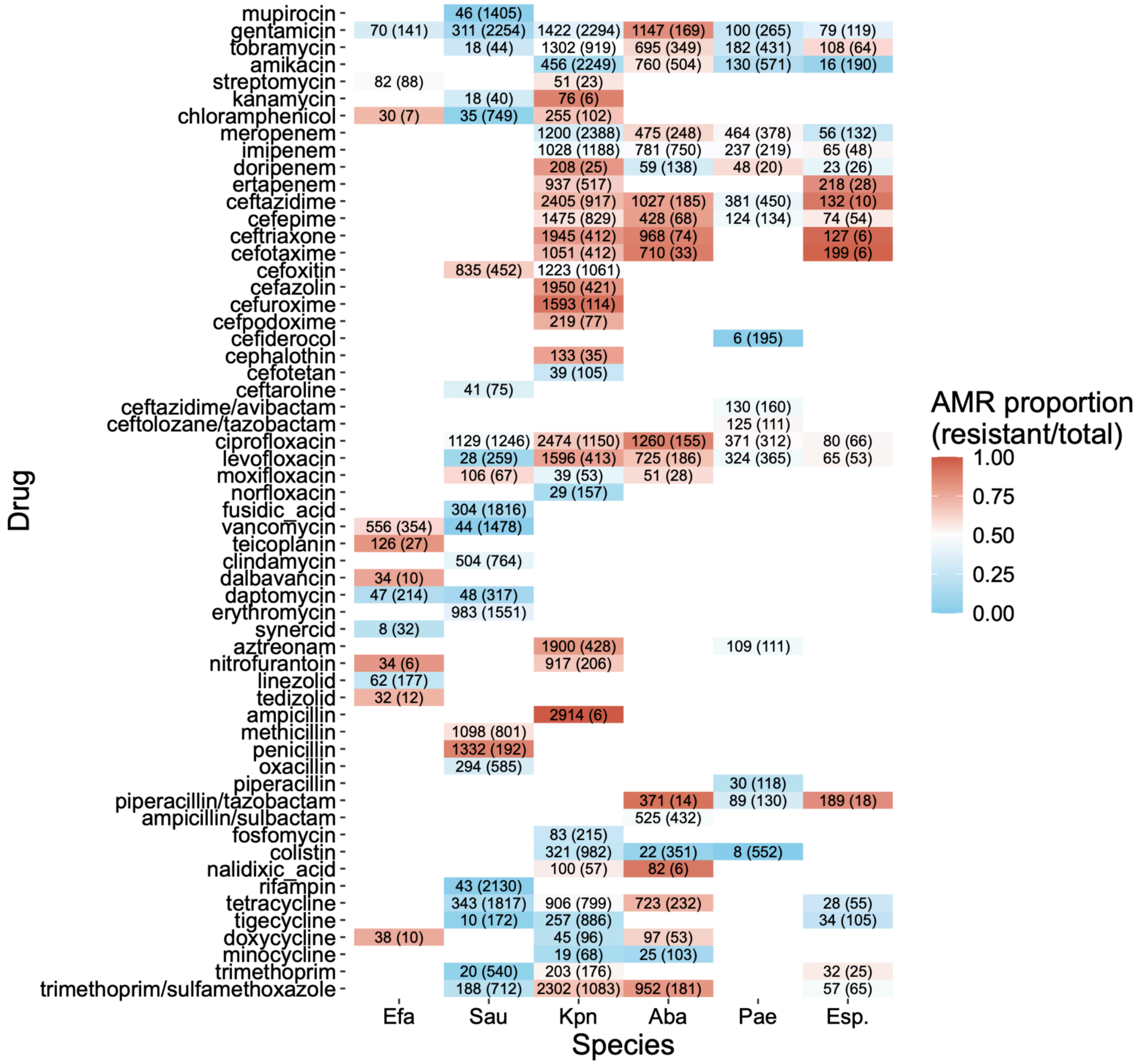


Figure S1. AMR prevalence across pathogens by drug class

Figure layout and colorscale as described in **Figure 2** legend, but here we report AMR proportions by individual drugs instead of drug class. Drugs are grouped by drug class and by data availability within each drug class (first by number of species, then by total number of isolates across species).

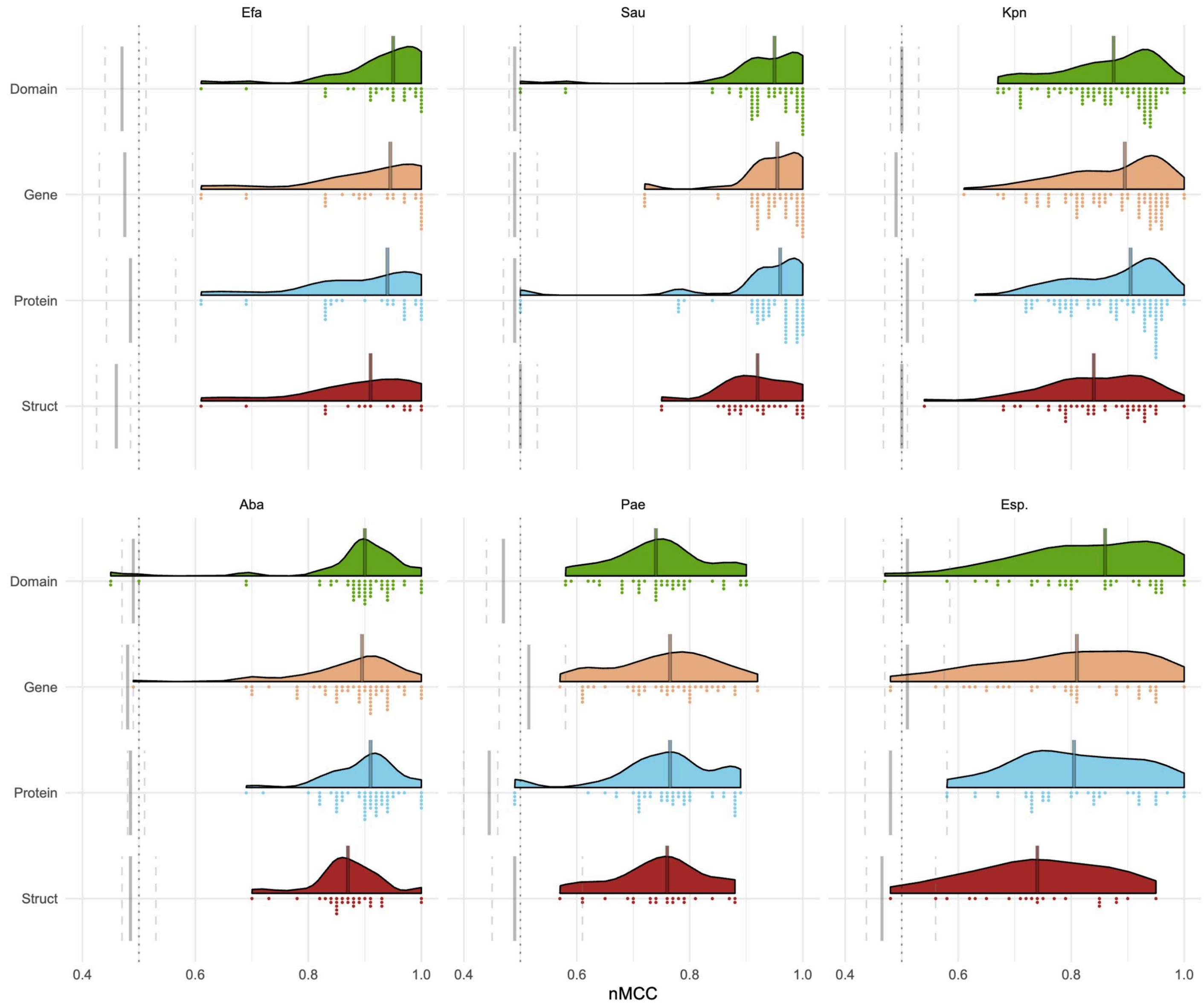


### Figure S2. Best performing feature scales across tested drugs per species.

Model performance is reported by nMCC (x-axis) for each scale (domains, genes, proteins, struct). Colored density distributions represent the spread of per-drug and data type models, which are individually represented by the colored points below, and the medians of the distribution are represented by solid lines. The solid and dashed grey lines represent the performance (nMCC) of the label shuffled baseline models' medians and interquartile ranges.

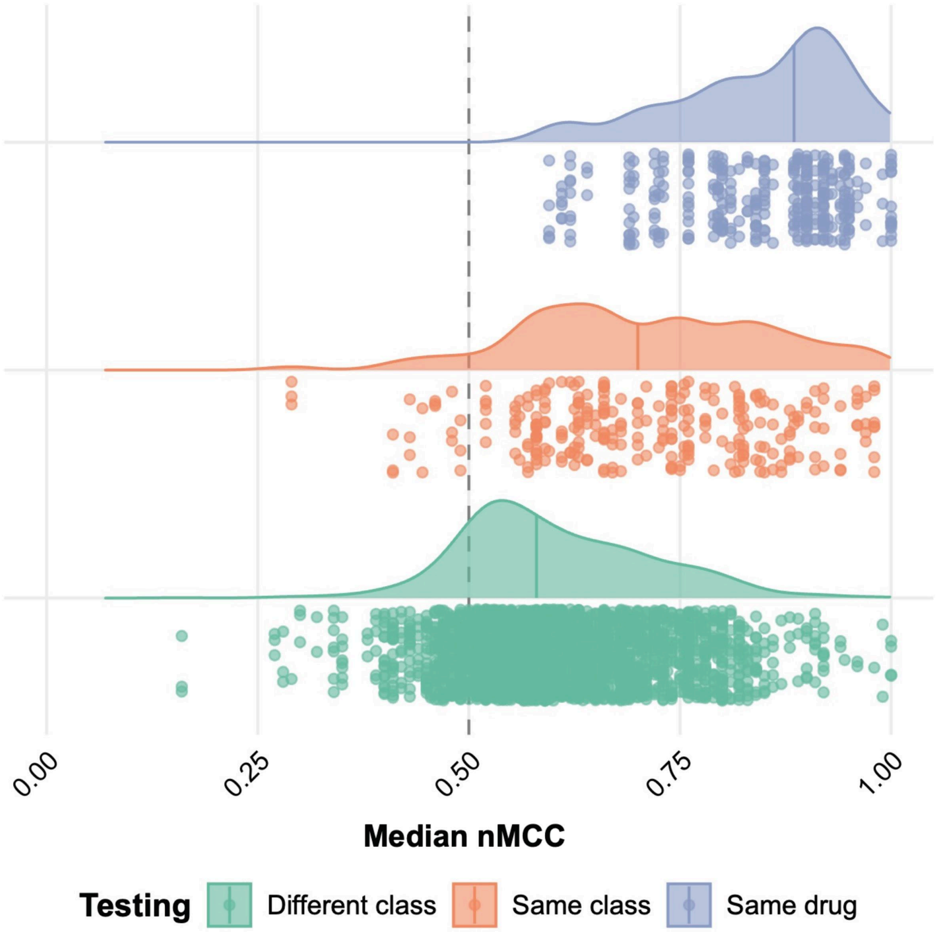


Figure S3. Cross-drug model performance across species, molecular scales, and drug classes.

Performance distribution of models trained on one drug and tested on others across ESKAPE. Each point represents the median nMCC calculated using the performance of all molecular scale and data type models for each tested drug. The plot shows that the models performed better when tested on the same drug (blue) or other drugs from the same class (orange) over other drug classes (green).

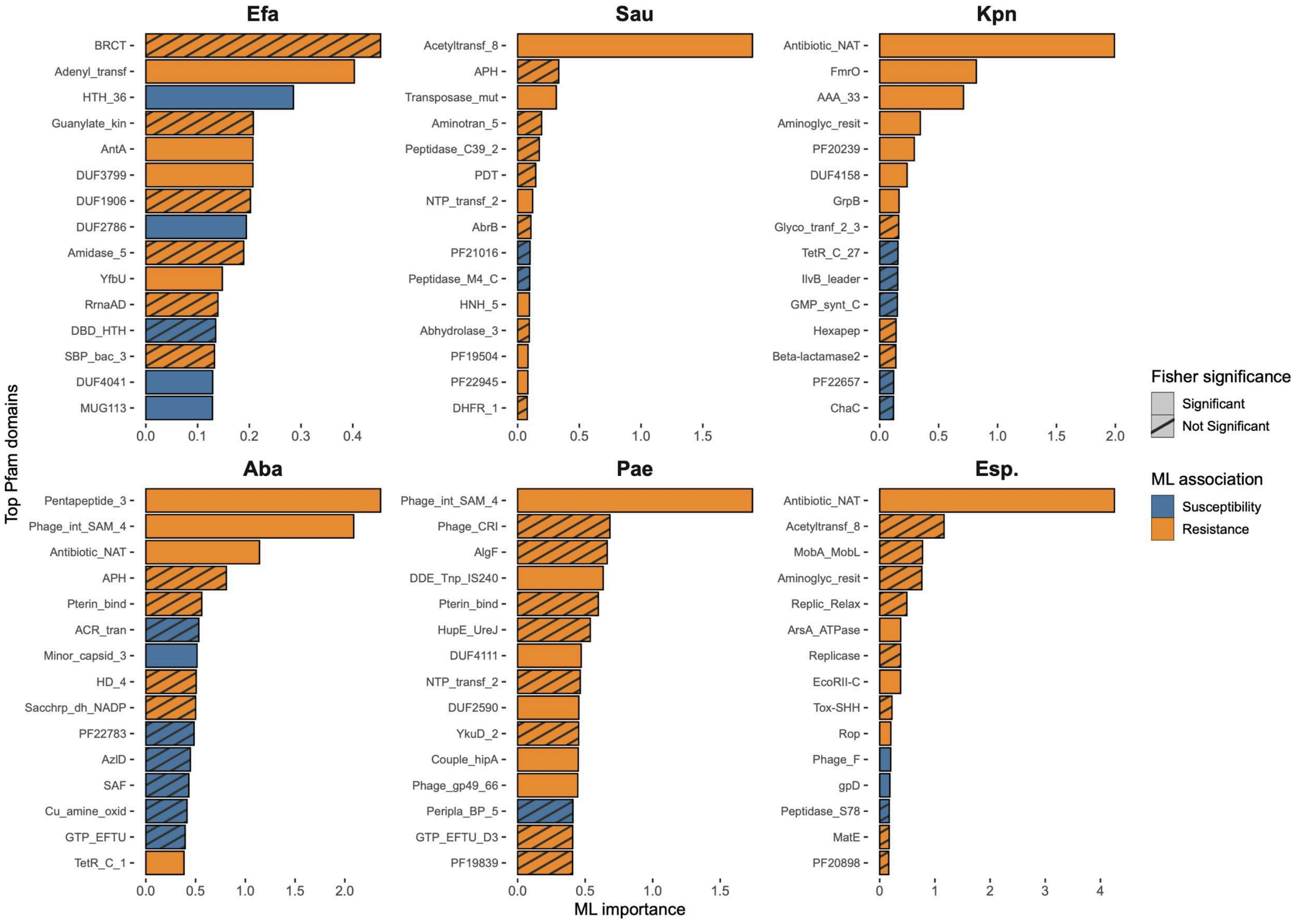


#### Figure S4. Demonstration of top ML features against Fisher's features.

We ran feature-wise two-sided Fisher's exact tests against AMR labels with Benjamini-Hochberg correction and recorded statistically significant features to compare against top predictors identified by our ML models. To illustrate, we show the features identified in our domain count gentamicin ML models across ESKAPE. Features associated with resistance and susceptibility are designated as orange and blue, respectively. ML features that were not statistically significant by Fisher's test are marked by bars with cross-stripes.

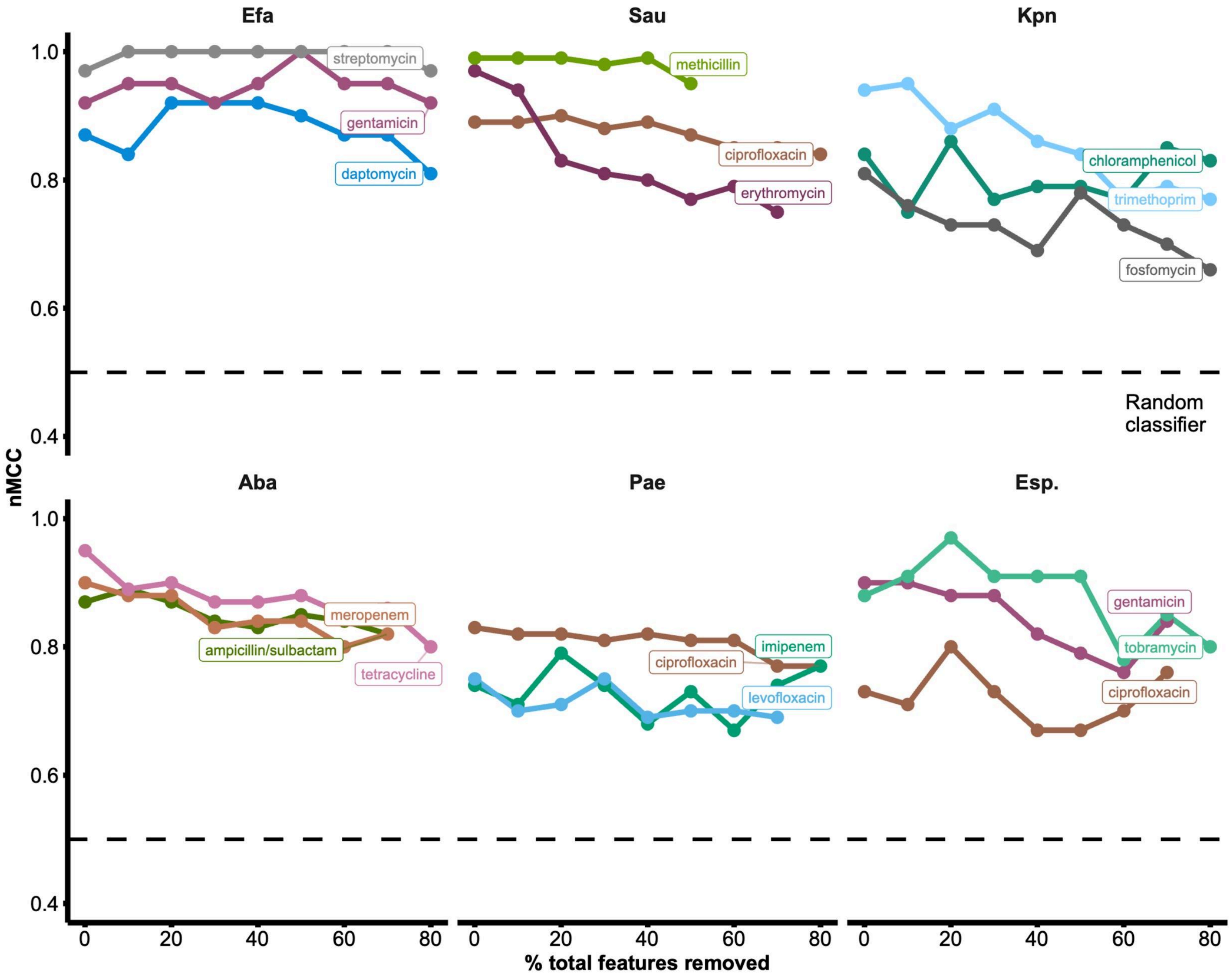


Figure S5. Removing top features iteratively with lasso regression.

To see how well models retain their predictive abilities when top features are removed, we started by training and testing with full datasets containing information about the presence/absence of genes. Once top features had been identified, we hid these from the models, retrained, retested, and repeated this process in increments corresponding to 10% of all genes in the respective datasets (relative to the original total number) per species-drug combination. In cases where all top features together comprised <10% of a dataset, genes were randomly selected without replacement for subsequent removal until 10% had been reached. We fixed the regularization parameter here to lasso regression to demonstrate its consolidation of the feature space to representatives from groups of genes with similar presence/absence patterns, allowing retention of AMR-associated genes across iterations. For illustrative purposes, we show only the three drugs with the most observations in their respective smallest AMR phenotype classes per species. This reduces the noise in our results by filtering out drugs with high iteration-to-iteration variability. Performance, as measured by nMCC, remains high with the removal of top features.

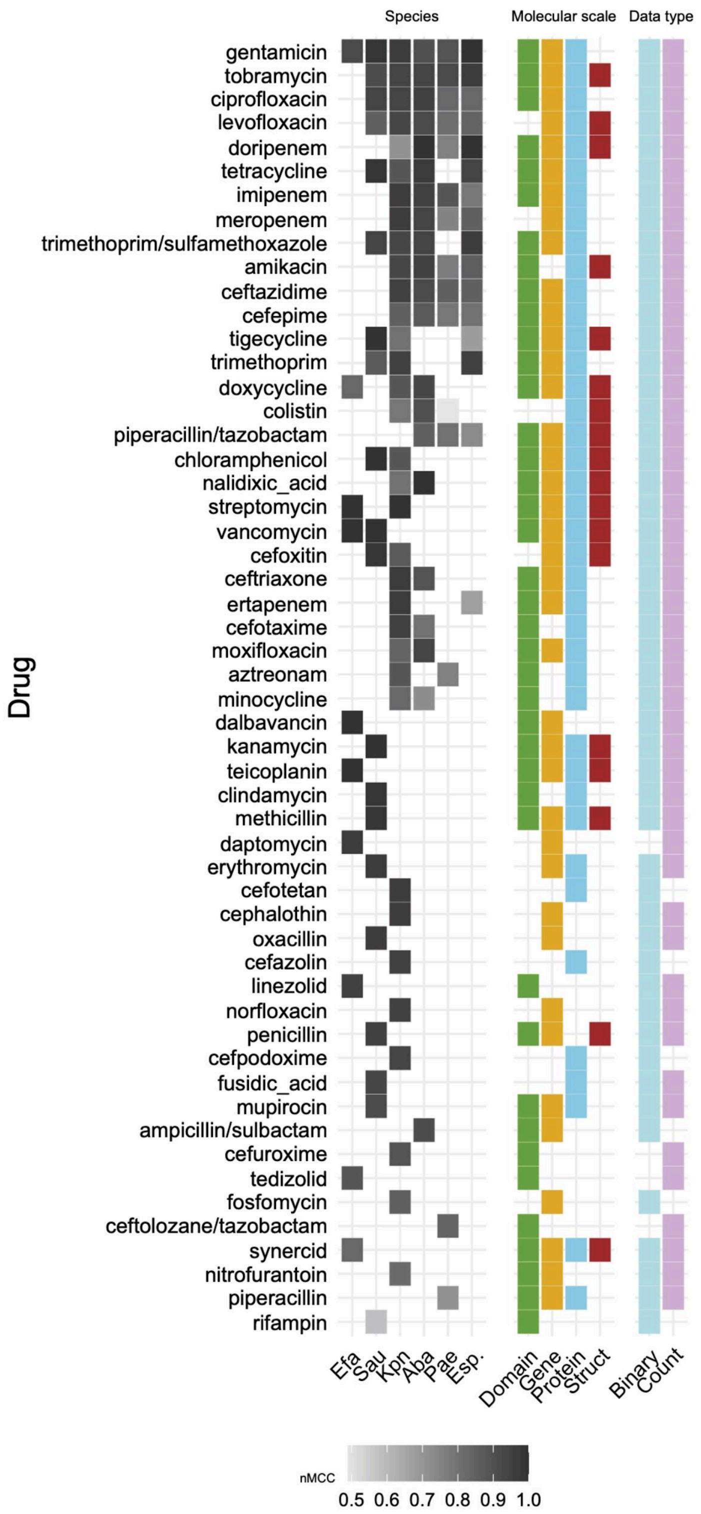


#### Figure S6. Best performing feature scales across tested drugs per species.

Grayscale colored tiles represent nMCCs for models, where black (1.0) shows perfect predictive performance, and light gray (0.5) indicates random performance. The drugs are ordered based on their performance and data availability across species. The next panels show tiles where that molecular scale or data type produced models with the top-ranked nMCCs. In cases where multiple molecular scales or data types are shown for a given drug class model, this indicates a tie in nMCC ranking. Feature scales and datatypes were comparable in performance, and overall model performance was high across most drugs and species.

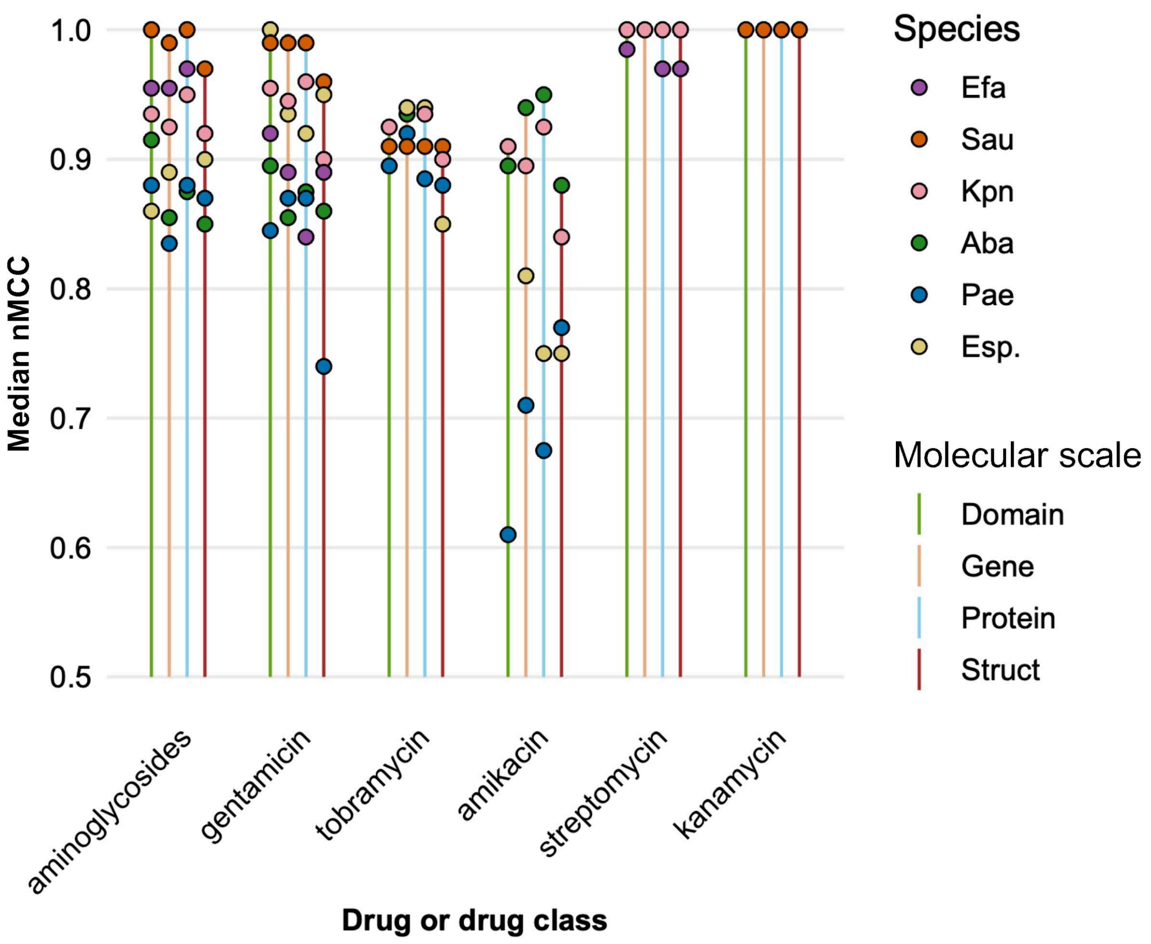


Figure S7. Aminoglycosides model performances across molecular scales for ESKAPE.

The median nMCC is calculated per molecular scale for each species and plotted as lollipops where the stems and heads are coloured by molecular scale and species, respectively; the height represents the median nMCC value.

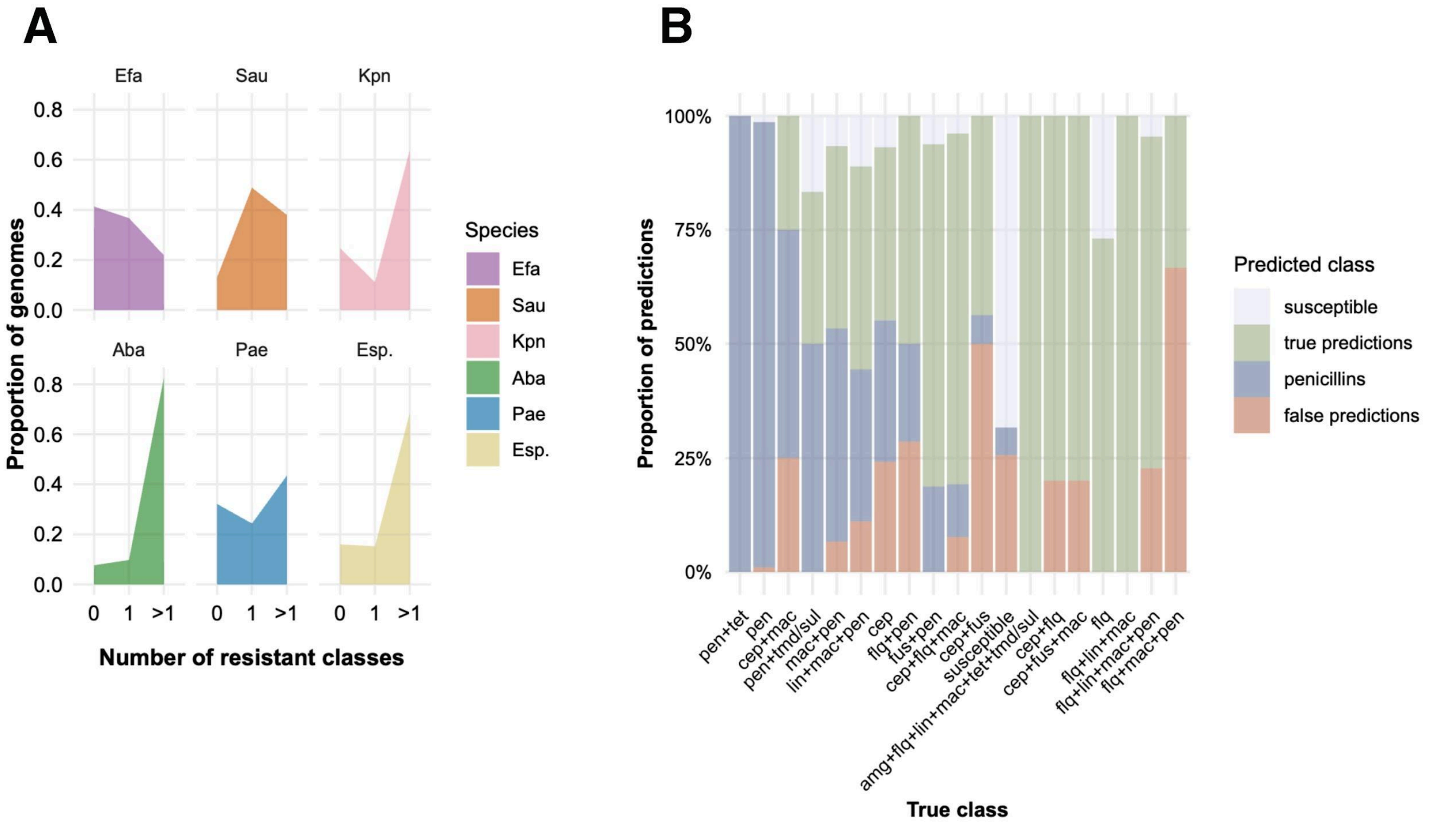


Figure S8. Proportions of predicted resistant classes across drug combinations.

**A. Fraction of ESKAPE genomes resistant to 0, 1, or multiple drug classes.** The number of classes each genome is resistant to is counted and grouped as 0 (susceptible), 1 (resistance to a single class), and >1 (resistance to multiple classes), and these relative frequencies are plotted per ESKAPE species. **B. The prediction bias in the *S. aureus* MDR multiclass model towards penicillins.** The multiclass model for drug resistance prediction in *S. aureus* genomes classified as susceptible, single drug resistant (flq, cep, or pen), or resistant to multiple classes (e.g., pen+tet). The colors in the stacked bar plot indicate the fraction of genomes per true class that are correctly ('true predictions') or incorrectly predicted ('false predictions'), and the fraction of times they are correctly/wrongly predicted as 'penicillins' or 'susceptible'. The abbreviations are explained in **Table S8**.

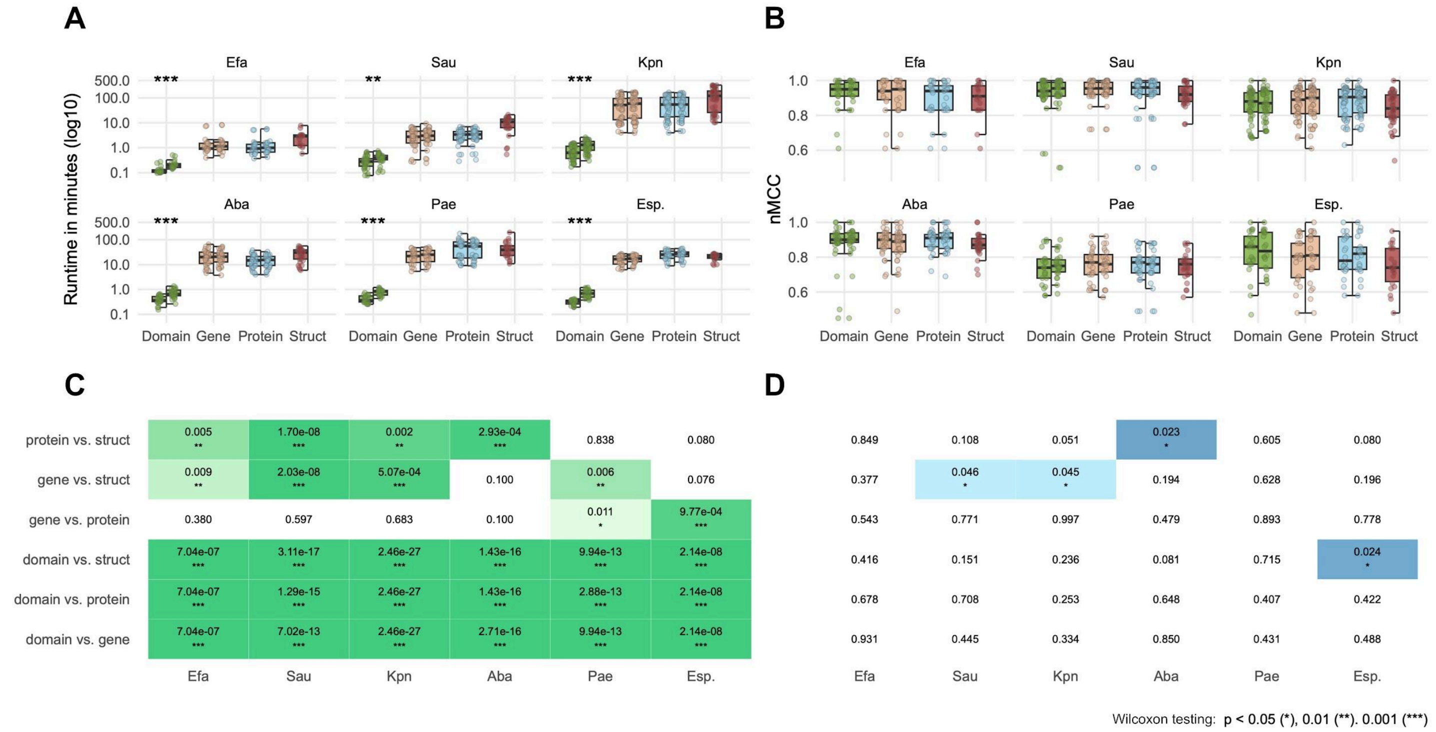


#### Figure S9. Model runtime comparisons across feature scales.

To compare another aspect of model performance beyond their predictive potential, we plotted per-model runtimes **(A)** in log-scaled minutes and nMCC **(B)** performances. Species across ESKAPE are faceted, molecular feature scales are plotted within each facet, and box plots are split to represent binary (left) and count (right) models for that feature scale. Structural variant features only exist as binary models. We performed Wilcoxon testing for pairwise comparison of runtimes and nMCCs between molecular scales across ESKAPE; the p-values and their level of significance are shown in panels **(C)** and **(D)**, which are visualized as color gradients.

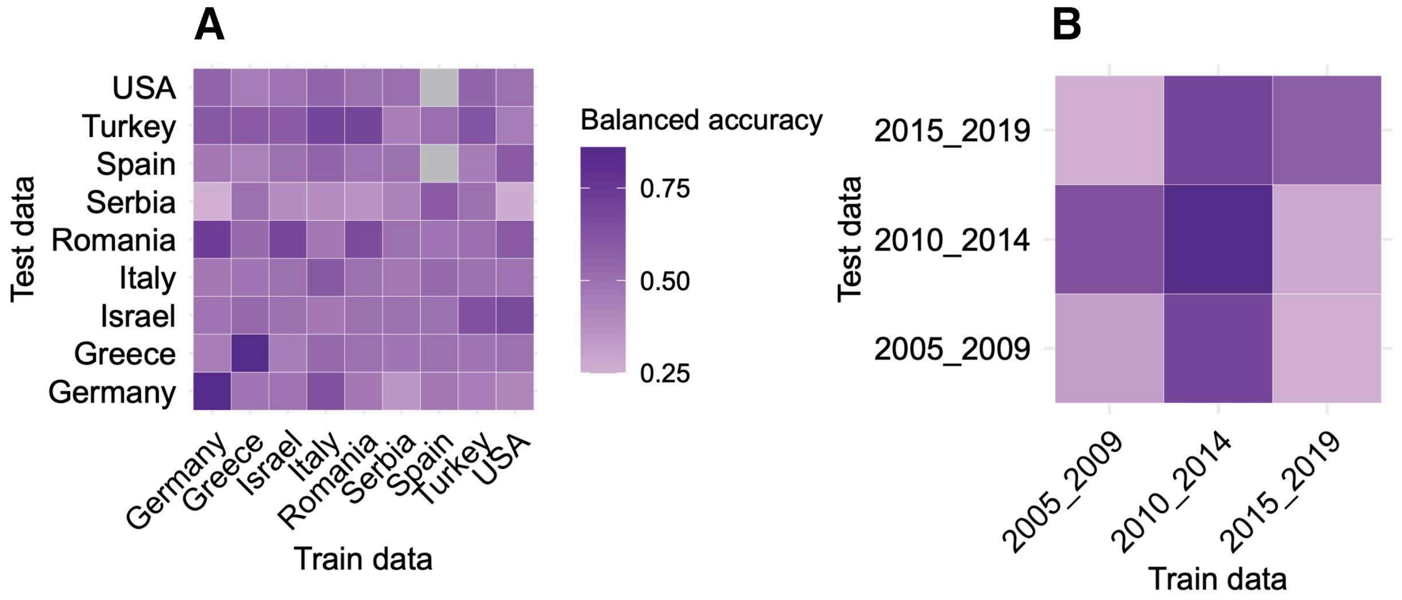


Figure S10. Geographical and temporal holdout testing performances.

**A. The performances of geographical holdouts for colistin models for *K. pneumoniae*.** For example, a model tested only on German data does best when tested on other German data, performance declines modestly when tested against Romanian data, and performance drops sharply when tested against Serbian data. **B. Temporal holdout testing for trimethoprim models in *K. pneumoniae*.** For example, models were trained on *K. pneumoniae* data isolated from the years 2005 to 2009, and tested against data from those years vs. later bins (e.g., 2015 to 2019). These holdout models show notable variations in performance when temporally limiting our models' data.

#### Supplementary tables

(The spreadsheet is attached separately.)

##### Table S1. Model performances across ESKAPE species, molecular scales, data types, drugs, and drug classes.

This table reports a summary of model data inputs and performance across drugs and drug classes among the ESKAPE pathogens. Per `species` for a given `drug` or `drug_class`, modeled per type of `data` and `scale`, the model performance (`nMCC`) is recorded, as well as the number of observations available in the data for that species and drug/drug class combination (`num_obs`) and the proportion of resistance seen in those observations (`res_prop`).

##### Table S2. Performances for shuffled baseline models.

As in **Table S1**, this table reports model performance and data summaries, but for the shuffled baseline control models. In these cases, the true phenotype labels (resistant/susceptible) were randomly shuffled in order to sever the connection between genotype (the genomic feature inputs) and phenotypes (the recorded AMR status).

##### Table S3. Performances for PCA models.

As in **Table S1**, this table reports model performance and data summaries, but for the models where feature spaces were reduced by principal components analysis (PCA).

##### Table S4. Performances for geographical holdout models.

As in **Table S1**, this table reports model performance and data summaries, but for the models subjected to geographic holdouts. For example, a model trained on tobramycin data from *K. pneumoniae* isolates from Ireland was tested on isolates from Australia (and other held-out isolates from Ireland; self-testing) to evaluate how such holdouts influence model performance, and how important features for predicting AMR differ from around the world.

##### Table S5. Performances for temporal holdout models.

As in **Table S1**, this table reports model performance and data summaries, but for the models subjected to temporal holdouts. That is, a model trained on tetracycline data from *S. aureus* isolates from the years 2000 to 2004 was tested on data isolated from the years 2015 to 2019 (as well as the training temporal bin; self-testing) to evaluate how model performance and top features would vary over time and as pathogens evolve, expand, and exchange genetic elements.

##### Table S6. Performances for cross-drug trained models.

As in **Table S1**, this table reports model performance and data summaries for the models trained on one drug, but tested against different drugs. That is, a model trained on doxycycline data from *K. pneumoniae* is tested for predicting minocycline resistance (different drugs within the same drug class), or a model trained on gentamicin data from *A. baumannii* is tested for predicting cefotaxime resistance (entirely different drug classes). This tests whether some features predictive of certain kinds of AMR may be generalizable or contribute to different types of resistance, which may be expected for same-class drugs, but may yield more interesting features if models are also predictive across classes.

##### Table S7. Shortened drug class names.

A table explaining the shortened drug class names used for brevity in some figures.

##### Table S8. Abbreviated drug class names.

A table explaining the abbreviated drug class names used for brevity in **Figure S8**.
